# Anti-GD2 antibody disrupts GD2:Siglec-7 interactions and synergizes with CD47 blockade to mediate tumor eradication

**DOI:** 10.1101/2021.03.19.436221

**Authors:** Johanna Theruvath, Marie Menard, Benjamin A.H. Smith, Miles H. Linde, Garry L. Coles, Wei Wu, Louise Kiru, Alberto Delaidelli, John L. Silberstein, Allison Banuelos, Shaurya Dhingra, Elena Sotillo, Sabine Heitzeneder, Aidan Tousley, John Lattin, Peng Xu, Jing Huang, Nicole Nasholm, Guillermo Nicolas Dalton, Andy He, Tracy C. Kuo, Emma R.B. Sangalang, Jaume Pons, Amira Barkal, Rachel Brewer, Kristopher D. Marjon, Payton L. Marshall, Ricardo Fernandes, Jennifer R. Cochran, Poul H. Sorensen, Heike E. Daldrup-Link, Irving L. Weissman, Julien Sage, Ravindra Majeti, Carolyn R. Bertozzi, William A. Weiss, Crystal L. Mackall, Robbie G. Majzner

## Abstract

The disialoganglioside GD2 is consistently overexpressed in neuroblastoma and osteosarcoma, and is variably expressed in other sarcomas, gliomas, neuroendocrine tumors, and epithelial cancers. Anti-GD2 antibodies have improved the survival rates of patients with neuroblastoma only when administered as part of intense chemotherapy-based cytotoxic regimens, which are associated with debilitating late effects including hearing loss, growth retardation, and secondary leukemias. Despite broad expression of GD2 on osteosarcoma, anti-GD2 antibody has not mediated significant antitumor activity in that disease or any other GD2+ cancers. CD47 is a checkpoint molecule overexpressed on tumor cells that inhibits macrophage activity, and CD47 blockade has demonstrated promising clinical activity in early human trials. We investigated whether anti-CD47 antibody could enhance the efficacy of anti-GD2 antibody in neuroblastoma and other GD2+ malignancies. We demonstrate substantial synergy of these two agents, resulting in the recruitment of tumor associated macrophages (TAMs) to mediate robust and durable anti-tumor responses. The responses are driven by GD2-specific factors that reorient the balance of macrophage activity towards phagocytosis of tumor cells, including disruption of a newly described GD2:Siglec-7 axis. These results demonstrate the unique synergy of combining anti-GD2 with anti-CD47, which has the potential to significantly enhance outcomes for children with neuroblastoma and osteosarcoma and will soon be investigated in a first-in-human clinical trial.

## Introductory Paragraph

The disialoganglioside GD2 is consistently overexpressed in neuroblastoma^1, 2^ and osteosarcoma^3–5^, and is variably expressed in other sarcomas^5^, gliomas^6^, neuroendocrine tumors^7^, and epithelial cancers^8^. Anti-GD2 antibodies have improved the survival rates of patients with neuroblastoma only when administered as part of intense chemotherapy-based cytotoxic regimens^9–12^, which are associated with debilitating late effects including hearing loss, growth retardation, and secondary leukemias^13–15^. Despite broad expression of GD2 on osteosarcoma, anti-GD2 antibody has not mediated significant antitumor activity in that disease or any other GD2+ cancers^16–18^. CD47 is a checkpoint molecule overexpressed on tumor cells that inhibits macrophage activity^19^, and CD47 blockade has demonstrated promising clinical activity in early human trials^20, 21^. We investigated whether anti-CD47 antibody could enhance the efficacy of anti-GD2 antibody in neuroblastoma and other GD2+ malignancies. We demonstrate substantial synergy of these two agents, resulting in the recruitment of tumor associated macrophages (TAMs) to mediate robust and durable anti-tumor responses. The responses are driven by GD2-specific factors that reorient the balance of macrophage activity towards phagocytosis of tumor cells, including disruption of a newly described GD2:Siglec-7 axis. These results demonstrate the unique synergy of combining anti-GD2 with anti-CD47, which has the potential to significantly enhance outcomes for children with neuroblastoma and osteosarcoma and will soon be investigated in a first-in-human clinical trial.

We measured phagocytosis of GD2+ neuroblastoma cells by human macrophages in vitro after treatment with anti-GD2 (dinutuximab, chimeric mouse/human IgG1 antibody), anti-CD47 (B6H12, murine IgG1 antibody), or a combination of both antibodies. The combination of anti-GD2 and anti-CD47 resulted in significant enhancement in phagocytosis of neuroblastoma cells (Figure 1a, b). *In vivo*, using an aggressive, orthotopic xenograft model of MYCN amplified neuroblastoma (KCNR), neither anti-GD2 antibody nor anti-CD47 antibody mediated significant anti-tumor effect, but the combination resulted in complete tumor eradication and long-term survival (Figure 1c-f). We also performed this experiment with the clinically validated humanized anti-CD47 antibody magrolimab (Hu5F9-G4, humanized IgG4 antibody)^21^, which demonstrated similarly profound anti-tumor activity when combined with anti-GD2 (Extended Data Figure 1a-c). To demonstrate the generalizability of our approach, we repeated this experimental setup with a MYCN non-amplified neuroblastoma cell line (CHLA255) in both a metastatic (Figure 1 g-j) and an orthotopic model (Extended Data Figure 1c-f). In both models, the combination of anti-GD2 and anti-CD47 mediated potent anti-tumor activity, resulting in cure of dual treated mice. If these results are validated in the clinic, this approach will provide a unique option for the treatment of neuroblastoma that does not require cytotoxic chemotherapy.

**Figure 1.**
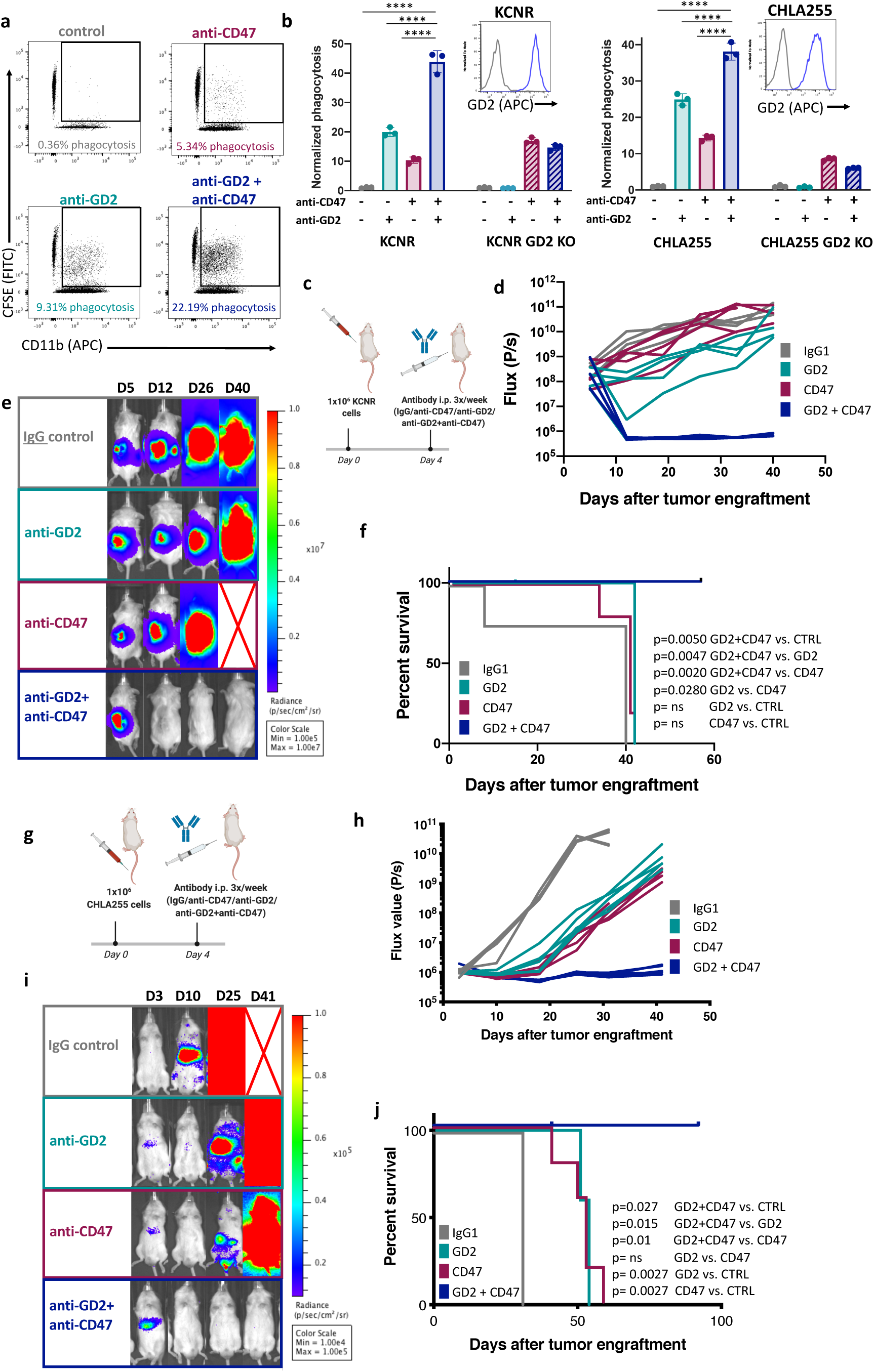
Anti-CD47 and anti-GD2 synergize to mediate significant anti-tumor activity in xenograft models of neuroblastoma. a. Representative flow-cytometry plots depicting the phagocytosis of KCNR cells labeled with CFSE co-cultured with human blood derived macrophages in the presence or absence of anti-GD2 mAb, anti-CD47 mAb or dual treatment. Plots are representative of at least three experiments each performed with three different blood donors. FITC, fluorescein isothiocyanate; APC, Allophycocyanin. **b**, Graphs show flow cytometry-based quantification of phagocytosis of KCNR and KCNR-GD2KO (left), CHLA255 and CHLA255-GD2KO (right) cell lines in the presence of anti-GD2 mAb, anti-CD47 mAb or dual treatment, compared with untreated control; results normalized to the phagocytosis in the untreated control for each cell line and blood donor. Error bars, s.d. of three experimental replicates. Statistical comparisons performed with one-way ANOVA with multiple comparisons correction, *P < 0.05, **P < 0.01, ***P < 0.001, ****P < 0.0001, ns P>0.05. Representative data from at least three experiments each performed with three different blood donors. Inset: flow histograms show levels of expression of GD2 on the surface of the neuroblastoma cell lines used for each assay, blue: Wildtype line, gray: GD2 KO (B4GALNT1 KO) line **c,** Experimental overview of the orthotopic neuroblastoma model: One million MYCN-amplified KCNR neuroblastoma cells expressing GFP-luciferase were implanted into the renal capsule. Four days later, mice were injected with IgG control, anti-GD2 mAb, anti-CD47 mAb or dual anti-GD2/anti-CD47 treatment every other day for three doses. **d,** Quantification of tumor progression for each individual mouse as measured by flux values acquired via bioluminescence (BLI) photometry. **e,** BLI images of representative mice from each treatment group shown in **d** at different time points. Red cross indicates deceased mouse. **f,** Survival curves for mice bearing tumors shown in **d**. Survival curves were compared using the log-rank test. **d-f** are representative of two independent experiments. N=5 (anti-GD2 mAb, anti-CD47 mAb, anti-GD2/anti-CD47), N=4 (IgG control). **g**, Experimental overview of the metastatic neuroblastoma model: One million MYCN non-amplified CHLA255 neuroblastoma cells expressing GFP-luciferase cells were engrafted into NSG mice by tail-vein injection. Four days later, mice were injected with IgG control, anti-GD2 mAb, anti-CD47 mAb or dual anti-GD2/anti-CD47 treatment every other day for three doses. **h,** Quantification of tumor progression for each individual mouse as measured by flux values acquired via BLI photometry. **i,** BLI images of representative mice from each treatment group shown in **h** at different time points. Red cross indicates deceased mouse. **j,** Survival curves for mice bearing tumors shown in **g**. Survival curves were compared using the log-rank test. **h-j** are representative of four independent experiments. N=5 mice per group in each experiment.

The immunocompromised NOD scid gamma (NSG) mice in our models lack lymphocytes involved in a coordinated anti-tumor immune response; it was therefore unclear how this therapeutic combination would perform in an immune competent model. Using allografts from the well described TH-MYCN syngeneic neuroblastoma model^22^, we investigated the addition of a CD47-blocker (ALX301) to a murine anti-GD2 antibody (14G2a, murine IgG2a) (Figure 2a). As the commercially available anti-murine CD47 antibodies are low affinity and minimally effective in vivo^23, 24^, we utilized a novel fusion of murine IgG1 to a mutated form of SIRPα capable of binding mCD47 with 3.41nM affinity (ALX301, Extended Data 2). The IgG1 component was engineered with an N297A mutation to minimize interaction with the murine Fc receptor, decreasing the potential for off-tumor toxicity. This molecule binds specifically to murine CD47 and is capable of blocking its interaction with SIRPα on murine macrophages.

**Figure 2.**
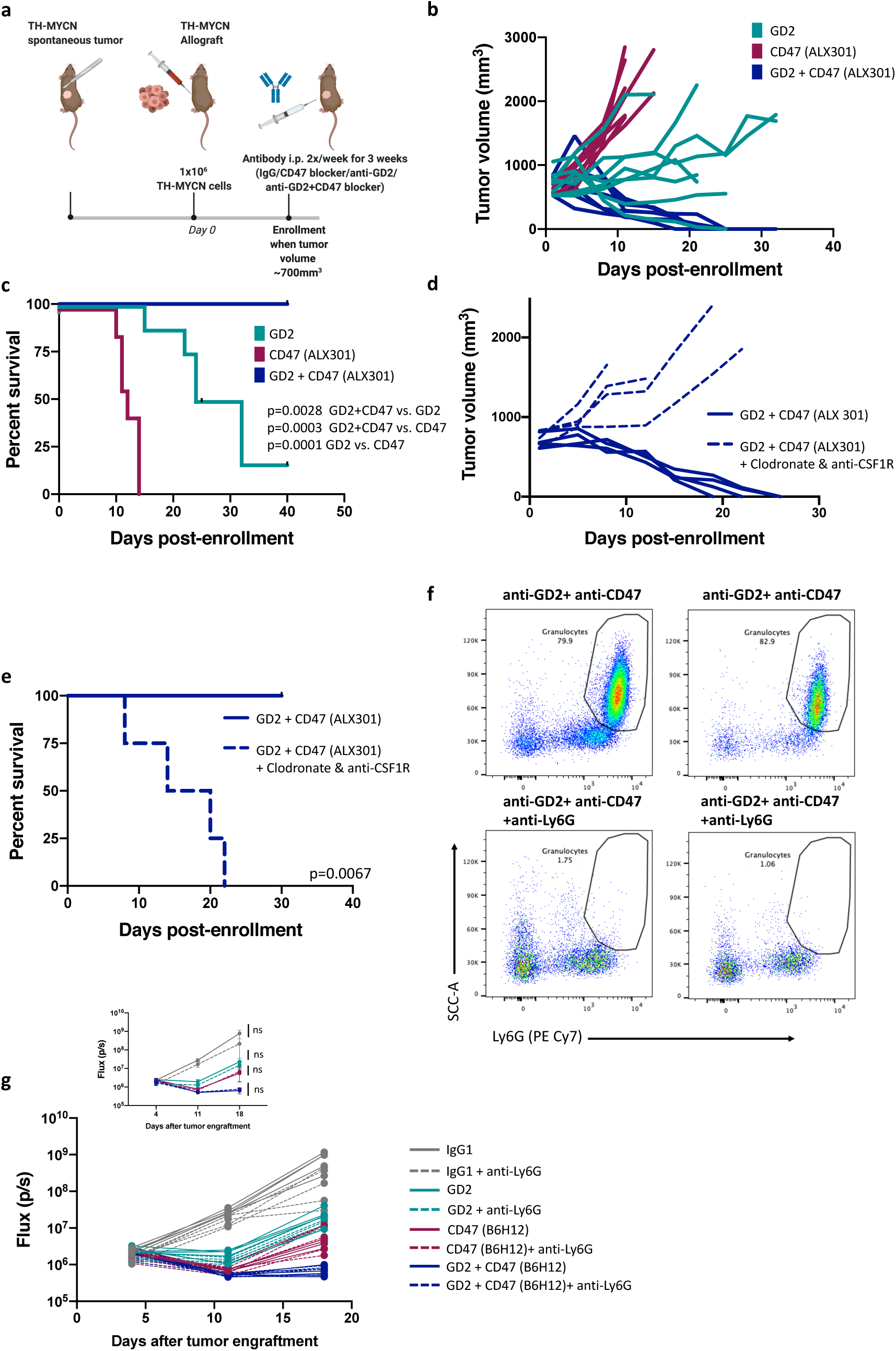
CD47 blockade licenses tumor associated macrophages (TAMs) to enhance the activity of anti-GD2 in a syngeneic model of neuroblastoma. **a,** Experimental overview of the syngeneic model of neuroblastoma: One million cells from de novo tumors from TH-MYCN mice were harvested and injected subcutaneously in the flank of female 129SvJ mice. Once tumors reached a volume of ∼700 mm3, mice were injected with IgG control, anti-GD2 mAb (14G2a), CD47 blocker (ALX301) or dual anti-GD2/CD47 blocker treatment 2 times/week for 3 weeks. b, Tumor progression for each individual mouse was followed using caliper measurements of tumor dimensions. **c,** Survival curves for mice bearing tumors shown in **a.** Survival curves were compared using the log-rank test. Data for **a-c** are from two pooled experiments. N=7 (anti-CD47, anti-GD2/anti-CD47), N=8 (anti-GD2) mice per group. **d,** Mice received clodronate liposomes and anti-CSF1R mAb or no depletion alongside anti-GD2/CD47 blocker. Tumor progression for each individual mouse was followed using caliper measurements of tumor dimensions. **e**, Survival curves for mice from **d.** Survival curves were compared using the log-rank test. N=4 mice per group. **f-g,** Mice treated with anti-Ly6G mAb were enrolled into the metastatic model of neuroblastoma as in Figure 1g. **f,** flow cytometry showing depletion of granulocytes in blood samples from mice treated with anti-Ly6G (bottom) vs control mice (top). **g,** Quantification of tumor progression for each individual mouse as well as mean values per treatment group (inset) as measured by flux values acquired via BLI photometry are shown. Statistical comparisons were made between Ly6G depleted and non-depleted groups using the repeated measures two way ANOVA (ns denotes not significant, P>0.05). N=5 mice per group.

As in the xenograft models, the addition of CD47 blockade to anti-GD2 dramatically enhanced anti-tumor responses, resulting in long-term, tumor-free survival of all combination-treated mice in the syngeneic model (Figure 2b-c). Myeloid cell depletion with clodronate and anti-CSF1R antibody ablated the anti-tumor efficacy (Figure 2d-e), indicating that, as in the NSG models, the response to anti-GD2/CD47 blocker is dependent on myeloid cells. To further delineate which myeloid cells were involved in the observed anti-tumor activity, we used anti-murine Ly6G antibody to selectively deplete neutrophils and granulocytes in NSG mice (Figure 2f), but observed no effect on the anti-tumor activity (Figure 2g), implicating macrophages/monocytes in the response to anti-GD2/anti-CD47.

Only those neuroblastoma patients with certain NK cell killer Ig-like receptor (KIR) and KIR-ligand genotypes derive benefit from anti-GD2 antibody therapy in the clinic^25, 26^, limiting its therapeutic reach. Although neuroblastoma tumors can be infiltrated by NK cells^27^ that can mediate anti-tumor activity, they are also highly infiltrated by macrophages^28^ which are thought to play an immune suppressive role^29^. In our models, when combined with anti-GD2 antibody, CD47 blockade reorients the myeloid compartment, rendering it capable of mediating significant anti-tumor activity which may be generalizable to patients independent of NK cells and KIR/KIR-L status.

Having demonstrated significant synergy of anti-GD2 and anti-CD47, we wondered whether CD47 blockade would similarly enhance the activity of other tumor specific antibodies. Both *in vitro* and *in vivo*, we evaluated for synergistic effects of anti-CD47 with an anti-B7-H3 antibody (enoblituzumab, a humanized Fc enhanced IgG1 antibody^30^). The addition of anti-B7-H3 to anti-CD47 was minimally effective in enhancing anti-tumor activity compared to anti-CD47 alone in either neuroblastoma or osteosarcoma xenografts (Extended Data Figure 3). We therefore hypothesized that GD2 specific factors were responsible for the enhanced activity observed in our xenograft and syngeneic models (Figures 1-2), which we then set out to define mechanistically.

We hypothesized that GD2 itself interacts with and inhibits immune cells in general, and macrophages in particular. GD2 is a disialoganglioside, containing two sialic acid residues. The Siglecs (Sialic acid-binding immunoglobulin-like lectins) are a family of 14 receptors expressed on immune cells that recognize and bind to specific sialic acid containing glycans and are capable of suppressing immune cell activity through their cytoplasmic immunoreceptor tyrosine-based inhibitory motif (ITIMs) domains^31^. We screened soluble versions of all commercially available human Siglecs for binding to the CHLA255 and KCNR neuroblastoma cell lines by flow cytometry and observed that Siglec-5, -7, -9, and -14 demonstrated different degrees of binding (Extended Data Figure 4a). Because tumor cells express numerous glycans capable of Siglec binding, we next sought to specifically assess whether GD2 binds any human Siglecs. To test this, we treated a GD2 negative cell line (Nalm6), with a cocktail of sialidases to remove all sialic acids and thus all Siglec ligands from the cell surface. We then incubated the sialidase treated cells with synthetic GD2, whose hydrophobic tail was readily integrated into the cell membrane (Figure 3a). We confirmed the presence of GD2 on the cell surface by flow cytometry (Extended Data Figure 4b) and then probed these GD2 coated cells with soluble human Siglecs. We discovered that GD2 specifically binds to Siglec-7 and to no other human Siglecs (Figure 3b). Additionally, lentiviral transfection of both B4GALNT1 (GD2 synthase) and ST8SIA1 (GD3 synthase) on Nalm6 to drive GD2 expression resulted in a substantial increase in Siglec-7 binding (Extended Data Figure 4c).

**Figure 3.**
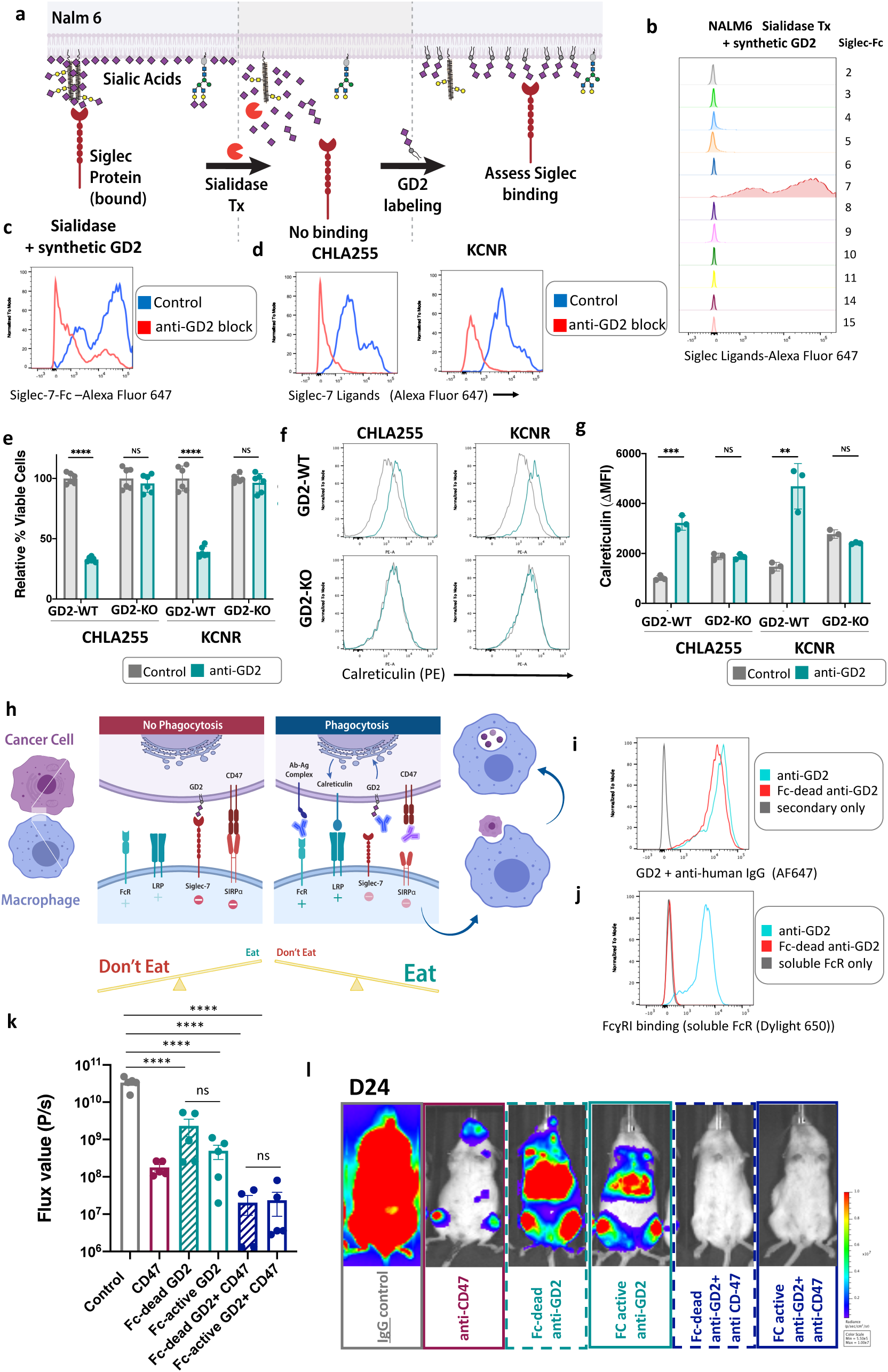
Anti-GD2 promotes tumor cell phagocytosis through blockade of Siglec-7 and upregulation of surface calreticulin. **a,** Experimental overview for the analysis of Siglec binding specificity to sialic-acid-stripped and GD2-coated Nalm6 cells. **b,** Flow cytometric histograms showing staining of Nalm6 cells pretreated with sialidase and coated with GD2 as in **a** with each individual recombinant Siglec. **c,** Flow cytometric analysis of Siglec-7 binding to Nalm6 treated as in **a** before (control, blue) and after incubation with anti-GD2 mAb (red). **d,** Flow cytometric analysis of Siglec-7 binding to neuroblastoma cell lines CHLA255 and KCNR before (control, blue) and after incubation with anti-GD2 mAb (red). **e,** Graph shows flow-based quantification of cell viability. Neuroblastoma cell lines and their isogenic GD2-KO versions were incubated with anti-GD2 mAb for 12 hours at 37 degrees and stained with DAPI. Percent of DAPI-populations were normalized to the untreated control for each cell line. Error bars, s.d. of 6 experimental replicates. Statistical comparisons performed with the unpaired t-test, *P < 0.05, **P < 0.01, ***P < 0.001, \*\*\*\**P* < 0.0001, ns P>0.05. **f,** Representative flow cytometric histograms of the expression of calreticulin on the surface of neuroblastoma cell lines (top) and their isogenic GD2-KO versions (bottom), after incubation for 12 hours at 37 degrees with anti-GD2 mAb. **g,** Graph shows flow cytometric quantification of the expression of calreticulin (ΔMFI) on the surface of neuroblastoma cell lines treated as in **e**. ΔMFI was calculated as the difference between the MFI in the PE channel of the calreticulin stained sample and the MFI in the PE channel of an isotype stained sample from the same experimental condition. Error bars, s.d. of 3 experimental replicates. Statistical comparisons performed as in **e. E-g** was performed at least three times. **h,** Proposed model of synergy of anti-GD2 and anti-CD47. At baseline, tumor cells express CD47 and GD2, which bind to SIRPα and Siglec-7 respectively on the surface of the macrophage, both of which are “Don’t Eat Me” signals that inhibit phagocytosis. Anti-GD2 blocks the GD2:Siglec-7 interaction and induces export of calreticulin from the endoplasmic reticulum to the cell surface of the tumor cell, upregulating an “Eat Me” signal. In the presence of anti-CD47, the balance of macrophage activity is shifted towards phagocytosis, resulting in synergistic anti-tumor activity. **i,** Flow cytometric analysis of anti-GD2 and Fc-dead anti-GD2 mAb binding to GD2 on CHLA255 cells. Both antibodies where detected with a fluorophore labeled secondary donkey anti-human IgG antibody. **j,** Flow cytometric detection of anti-GD2, but not Fc-dead anti-GD2, bound to GD2 on CHLA255 cells detected with soluble Fc*γ*RI conjugated with Dylight 650. **k-l,** Metastatic neuroblastoma model as described in Figure 1g. Mice were injected with IgG control, Fc-active anti-GD2 mAb, Fc-dead anti-GD2 mAb, anti-CD47 mAb, dual anti-CD47/Fc-active anti-GD2, or dual anti-CD47/Fc-dead treatment for two doses. **k,** Quantification of tumor burden for individual mice on Day 24 as measured by flux values acquired via bioluminescence (BLI) photometry. **l,** BLI images of representative mice from each treatment group shown in **l.** Error bars, s.e.m, N=5 mice per group, Statistical comparisons performed with one-way ANOVA with multiple comparisons correction, *P < 0.05, **P < 0.01, ***P < 0.001, \*\*\*\**P* < 0.0001, ns P>0.05.

Siglec-7 is expressed on human macrophages and NK cells, consistent with the immune infiltrate common in neuroblastoma^27, 28^, and is capable of suppressing immune cell activity through its cytoplasmic ITIM domain^32, 33^. Siglec-7:GD2 binding on Nalm6 cells artificially coated with GD2 was also interrupted by blocking with an anti-GD2 antibody (Figure 3c). Additionally, we found robust binding of Siglec-7 to neuroblastoma cell lines that was specifically disrupted by anti-GD2 antibody (Figure 3d), demonstrating that anti-GD2 antibodies are capable of blocking binding to Siglec-7, potentially preventing its suppressive effects on the immune system. These data define the interaction of GD2 with Siglec-7 as a novel axis that can be disrupted therapeutically by anti-GD2 antibodies. Similarly, the glycoprotein CD24 expressed on tumor cells was recently shown to bind to Siglec-10, also expressed by macrophages, inhibiting phagocytosis^34^. These data indicate an important role for Siglec molecules in governing phagocytosis and anti-tumor immunity based on the surface glycome of tumor cells.

Previous reports have found that GD2 ligation can result in cell death in neuroblastoma and melanoma cells in the absence of immune effector cells^35, 36^. We confirmed these findings with multiple neuroblastoma cell lines, demonstrating that >50% of tumor cells are killed in response to GD2 ligation *in vitro* (Figure 3e, Extended Data Figure 5). The cells that remain are not unaffected; we found increased levels of surface calreticulin (Figure 3f-g, Extended Data Figure 5), a pro-phagocytic “Eat Me” signal that is shuttled to the surface from the endoplasmic reticulum in response to cell stress^37^. Therefore, GD2 ligation by anti-GD2 antibody primes cells for removal by the immune system. Taken together, these two findings explain the strong synergy of anti-GD2 and anti-CD47. We propose a model wherein blocking GD2 binding to Siglec-7 removes an inhibitory “Don’t Eat Me” signal from the tumor to the macrophages, while GD2 ligation results in upregulation of an “Eat Me” signal on the tumor cell. In the presence of anti-CD47, the balance of macrophage activity is shifted towards phagocytosis, resulting in potent anti-tumor activity (Figure 3h).

To demonstrate the functional consequences of GD2 blocking and ligation, we generated an Fc-dead version of dinutuximab^38^. This antibody is capable of binding GD2 on tumor cells (Figure 3i) but does not bind human Fc receptor (Figure 3j) nor does it fix complement. Despite its inability to directly interact with macrophages, the antibody significantly synergized with anti-CD47 antibody *in vivo*, demonstrating similar activity to the Fc-live version of the antibody when combined with CD47 blockade (Figure 3k-l). Thus, rather than simply providing an activating Fc domain to macrophages, treatment with anti-GD2 antibody significantly tilts the balance of macrophage activity towards phagocytosis, resulting in profound anti-tumor activity when combined with CD47 blockade.

GD2 is also expressed on other malignancies, including the pediatric bone tumor osteosarcoma^3–5^. However, anti-GD2 antibody has not mediated significant clinical benefit for most patients with osteosarcoma^16^. There have been no major advances in the treatment of osteosarcoma for decades and patients with metastases or relapse suffer extremely poor outcomes^39^. Virtually all patients with osteosarcoma receive the same chemotherapy regimens as thirty years ago, which are associated with the highest frequency of late effects of any pediatric cancer^40^. Many osteosarcoma tumors are infiltrated by macrophages^41^, so we hypothesized that the addition of anti-CD47 would license these TAMs for anti-tumor activity in the setting of anti-GD2.

We confirmed that the combination of anti-GD2 and anti-CD47 was effective *in vitro* using phagocytosis assays with osteosarcoma cell lines (Figure 4a). In an orthotopic xenograft model of osteosarcoma (MG63.3) (Figure 4b), we observed no effect of either single antibody treatment, but strong activity of the combination anti-GD2/anti-CD47, with significantly delayed tumor growth (Figure 4c) and prolonged survival (Figure 4d). Combination treatment was associated with an increase in macrophage infiltration and expression of inducible nitric oxide synthase (iNOS), consistent with the licensing of TAMs to mediate anti-tumor activity (Figure 4e-h). In parallel, by flow cytometry, we observed an increase in macrophage infiltration in combination treated mice (Extended Data Figure 6a), with a decreased percentage of immunosuppressive, M2 macrophages (Figure 4i, Extended Data Figure 6b).

**Figure 4.**
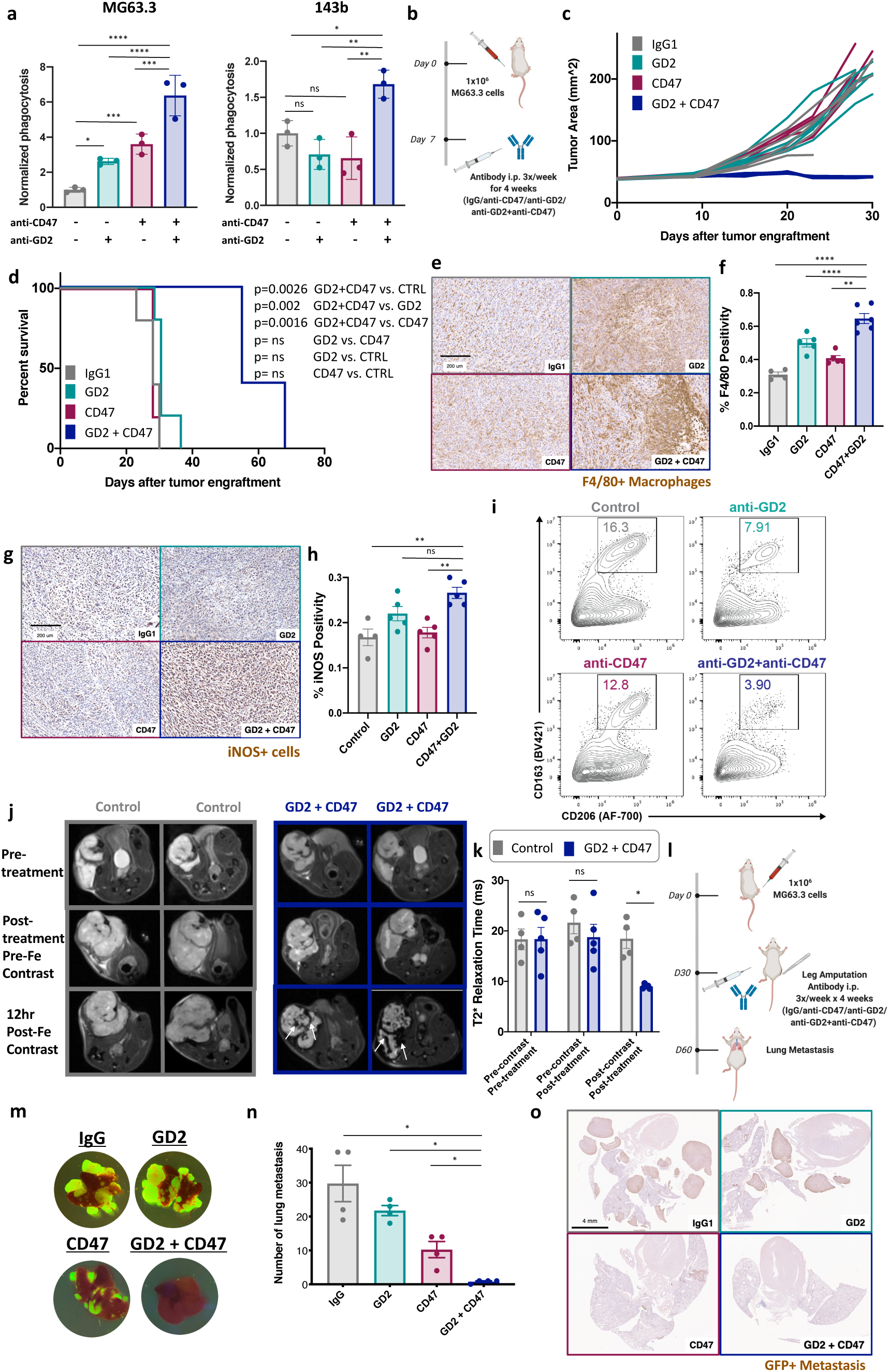
Anti-CD47 and anti-GD2 unleash TAM activity in osteosarcoma, resulting in meaningful anti-tumor responses. **a,** Graphs show flow cytometry-based quantification of phagocytosis of MG63.3 (left) and 143b (right) osteosarcoma cell lines co-cultured with human blood derived macrophages in the presence of anti-GD2 mAb, anti-CD47 mAb or dual treatment, compared with untreated control; results normalized to the phagocytosis in the untreated control for each cell line and blood donor. Error bars, s.d. of three experimental replicates. Statistical comparisons performed with one-way ANOVA with multiple comparisons correction, *P < 0.05, **P < 0.01, ***P < 0.001, ****P < 0.0001, ns P>0.05. Representative data from at least three experiments each performed with three different blood donors. **b,** Experimental overview of the orthotopic model of osteosarcoma: Periosteal injection of one million MG63.3 osteosarcoma cells expressing GFP. Seven days later, mice were injected with IgG control, anti-GD2 mAb, anti-CD47 mAb or dual anti-GD2/anti-CD47 treatment 3 times/week for 4 weeks. **c,** Tumor progression for each individual mouse was followed using caliper measurements of tumor dimensions. **d**, Survival curves for mice bearing tumors shown in **c.** Survival curves were compared using the log-rank test. **b-d** are representative of three independent experiments. N=5 mice per group. **e-i** Mice were treated as in **b** but tumors were allowed to grow for 21 days before initiation of treatment with IgG control, anti-GD2 mAb, anti-CD47 mAb or dual anti-GD2/anti-CD47. After one week of treatment, tumors were harvested for immunohistochemistry (IHC) and flow cytometric analysis. **e,** Representative IHC images showing detection of macrophages via staining with anti-F4/80 on tumors harvested from mice treated with indicated mAbs. **f**, Quantification of percent of positive F4/80 staining obtained from IHC analysis of 4-5 tumors from each treatment group. **g,** Representative IHC images showing staining with anti-iNOS on tumors harvested from mice treated with indicated mAbs. **h**, Quantification of percent of positive iNOS staining obtained from IHC analysis of 4-5 tumors from each treatment group**. E-h** Error bars, s.e.m. of 4-5 biologic replicates. Statistical comparisons were made as in **a**. **i,** Representative flow plots of M2 macrophages as defined by CD163+ and CD206+, gated on CD45+, CD11b+, F4/80 + macrophages. **j,** Representative magnetic resonance images (MRI) of mice bearing three-week-old tumors (top row), same mice one week after receiving control or anti-CD47/anti-GD2 combination treatment before (middle row) or 12 hours after (bottom row) injection of ferumoxytol contrast. Arrows indicate hypointense (dark) areas indicating iron-labeled macrophages in tumor tissue. **k,** Quantification of T2* relaxation time of tumors of mice treated as in **j.** Error bars, s.e.m. of 4 biologic replicates. Statistical comparisons were made between groups with the Mann-Whitney Test, p values as in **a**. **l,** Experimental overview of the metastatic model of osteosarcoma: Orthotopic MG63.3-GFP derived tumors were allowed to grow until most tumors were 10-12 mm in one direction before the mouse underwent leg amputation followed by administration of IgG control, anti-GD2 mAb, anti-CD47 mAb or dual anti-GD2/anti-CD47 3 times/week for 4 weeks. Lungs were harvested 30 days after amputation. **m,** Representative images of macroscopic analysis of GFP+ metastases in lungs from mice treated with indicated antibodies. **n**, Quantification of number of metastases per mouse for each treatment group. Error bars, s.e.m. of *n* = 4 biologic replicates. Statistical comparisons were made between groups as in **a**. Representative of two independent experiments. **o,** Representative IHC images showing detection of lung metastases via staining with anti-GFP on tumors harvested from mice treated with indicated mAbs.

We have previously shown that tumor macrophage infiltration can be measured by magnetic resonance imaging (MRI) after intravenous administration of ferumoxytol nanoparticles that are selectively phagocytosed by macrophages^42^. We imaged a cohort of control and anti-GD2/anti-CD47 treated mice with osteosarcoma xenografts by MRI at twelve hours after intravenous administration of ferumoxytol. Compared with control mice, we observed significantly reduced T2* relaxation values in combination treated mice, indicative of increased ferumoxytol retention in the tumor tissue and consistent with enhanced macrophage infiltration, validating this approach for use as a potential biomarker in patients that would not require surgical biopsies, which are often not feasible in pediatric clinical trials (Figure 4j-k).

The majority of mortality from osteosarcoma is driven by lung metastases, which are present but undetectable at the time of initial tumor diagnosis and treatment^39^. In a recent clinical trial of children and young adults with relapsed osteosarcoma who were rendered disease free by surgery, adjuvant treatment with dinutuximab (anti-GD2 antibody) and granulocyte-macrophage colony-stimulating factor did not prevent further relapses^16^. Drawing on an established model of pulmonary metastatic osteosarcoma^43^, we treated mice post-amputation with a control antibody, anti-GD2, anti-CD47, or a combination of anti-GD2/anti-CD47 (Figure 4l). Anti-GD2 treatment alone did not alter the number or size of metastases, in line with findings from the recent clinical trial^16^. While single agent anti-CD47 reduced the number and size of metastases, only the combination of anti-GD2 and anti-CD47 eradicated nearly all metastases (Figure 4m-o). Therefore, this approach may represent a novel mechanism to prevent pulmonary relapse in osteosarcoma, an area of pressing clinical need.

GD2 is also expressed on small cell lung cancer (SCLC)^7, 44^, an adult malignancy with especially poor outcomes despite adoption of immunotherapy into frontline regimens^45, 46^. A recent randomized trial of chemotherapy with or without dinutuximab failed to show any advantage to treatment with anti-GD2 antibody in patients with SCLC^17^. To evaluate whether anti-GD2/anti-CD47 could unleash TAMs in an additional GD2+ malignancy, we tested the combination in a xenograft model of SCLC and observed significantly enhanced anti-tumor activity and survival (Extended Data Figure 7). Therefore, combination with anti-CD47 may expand the clinical use of GD2-targeting antibodies to other GD2+ diseases beyond neuroblastoma.

In conclusion, we have demonstrated synergy of anti-GD2 and anti-CD47 in seven separate murine models in multiple histologies, offering a novel, chemotherapy free regimen to treat GD2+ malignancies. Anti-GD2/anti-CD47 recruits TAMs into the anti-tumor response, rendering them capable of mediating complete tumor clearance. The significant anti-tumor activity in our models derives from two GD2 specific factors. GD2 ligation directly results in upregulation of surface calreticulin on tumor cells, driving macrophage phagocytosis. Like dinutuximab, the anti-CD20 antibody rituximab is known to drive direct tumor cell killing in lymphoma^47, 48^, and this may similarly account for the enhanced activity observed in human trials when combined with CD47 blockade^21^. Second, we have defined the ligand for GD2 to be Siglec-7 and demonstrated a role for anti-GD2 antibodies in blocking this interaction to unleash immune cell activity. Blocking Siglec binding to glycolipids and glycopeptides is an emerging field of interest^49, 50^, and agents that target Siglec molecules have demonstrated early activity in the clinic^51^. This work lays a new framework for defining which types of tumor specific antibodies are best combined with CD47 blockade, namely those capable of further enhancing phagocytosis by blocking other “Don’t Eat Me” signals and/or increasing expression of “Eat Me” signals. Our novel, mechanistically driven combination of anti-GD2/anti-CD47 has the potential to extend the benefits of dinutuximab without the use of chemotherapy to patients with neuroblastoma and to reach patients with osteosarcoma and other GD2+ malignancies.

## Material & Methods

### Cells and culture conditions

Human cell lines used in these studies were supplied by the following sources: MG63.3 and 143b by C. Khanna (NCI, NIH, Bethesda, MD), Nalm6 (ATCC), KCNR and CHLA255 by R. Seeger (Keck School of Medicine, USC), and NCI-H69 SCLC (ATCC). Some cell lines were stably transduced with GFP and firefly luciferase using retrovirus as previously described^52^. GD2 knockout was achieved in KCNR and CHLA255 tumor cells using CRISPR-Cas9 to target the gene encoding GD2 synthase (B4GALNT1) and then FACS sorting as previously described for other cell lines^6^. Nalm6 was lentivirally transduced with B4GALNT1 and ST8SIA1 as previously described^53^. All tumor cell lines were cultured in RPMI-1640, supplemented with 10% heat-inactivated FBS (Gibco, Life Technologies), 10mM HEPES, 100U/mL penicillin, 100 μg/ml streptomycin and 2mM L-glutamine (Gibco, Life technologies).

### Flow Cytometry

Data was collected with an LSR Fortessa X-20, BD LSR II (both BD Bioscience) or Cytek Aurora Cytometer (Cytek) and analyzed using FlowJo software. Cells were harvested, washed with FACS buffer (PBS supplemented with 2% FBS) and stained for 30 minutes in the dark on ice. Cells were washed with FACS buffer after each incubation step. Cells were gated on viable cells and singlet discrimination (FSC-A/FSC-H) was performed before further analysis.

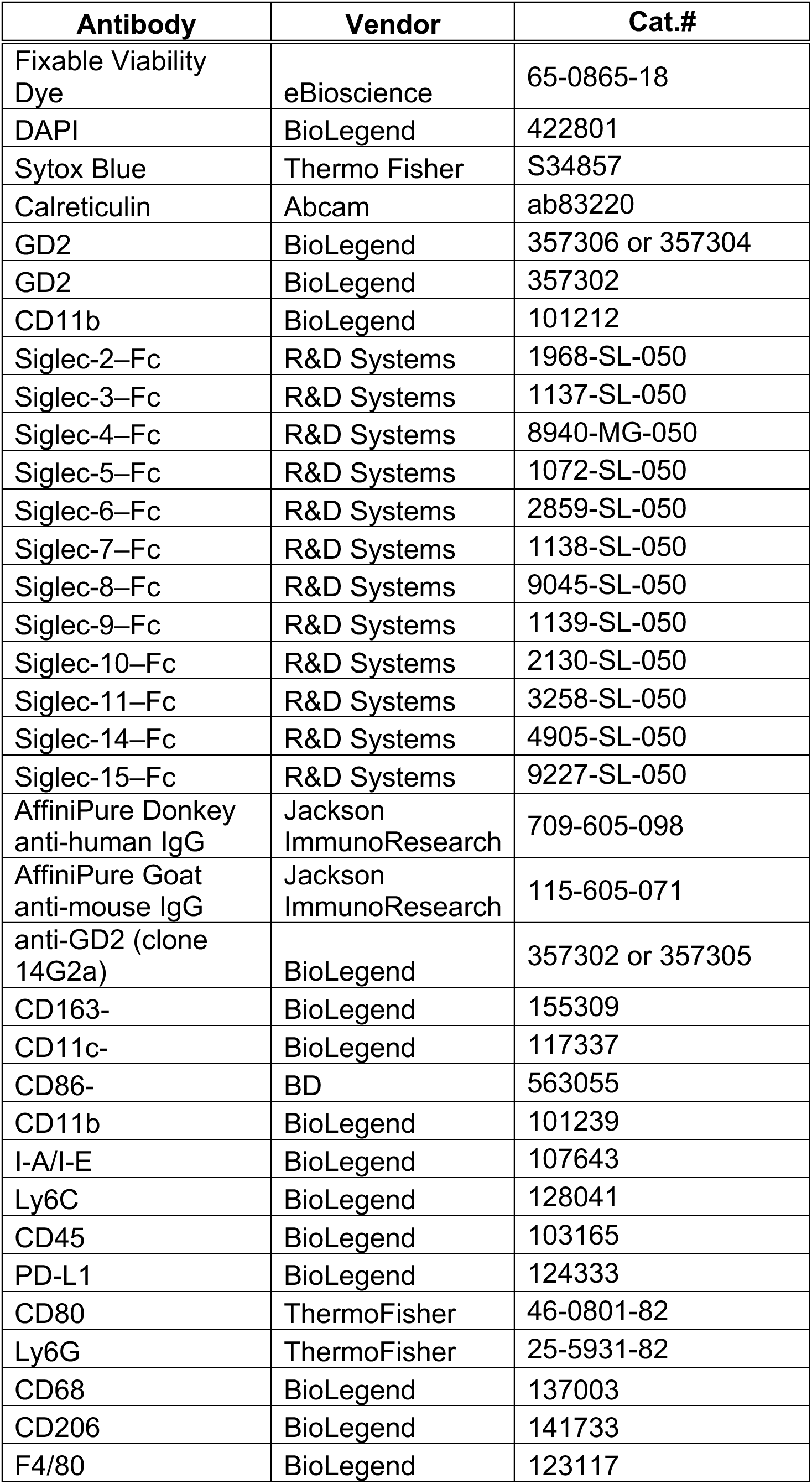

| Antibody | Vendor | Cat.# |
| --- | --- | --- |
| Fixable Viability Dye | eBioscience | 65-0865-18 |
| DAPI | BioLegend | 422801 |
| Sytox Blue | Thermo Fisher | S34857 |
| Calreticulin | Abcam | ab83220 |
| GD2 | BioLegend | 357306 or 357304 |
| GD2 | BioLegend | 357302 |
| CD11b | BioLegend | 101212 |
| Siglec-2–Fc | R&D Systems | 1968-SL-050 |
| Siglec-3–Fc | R&D Systems | 1137-SL-050 |
| Siglec-4–Fc | R&D Systems | 8940-MG-050 |
| Siglec-5–Fc | R&D Systems | 1072-SL-050 |
| Siglec-6–Fc | R&D Systems | 2859-SL-050 |
| Siglec-7–Fc | R&D Systems | 1138-SL-050 |
| Siglec-8–Fc | R&D Systems | 9045-SL-050 |
| Siglec-9–Fc | R&D Systems | 1139-SL-050 |
| Siglec-10–Fc | R&D Systems | 2130-SL-050 |
| Siglec-11–Fc | R&D Systems | 3258-SL-050 |
| Siglec-14–Fc | R&D Systems | 4905-SL-050 |
| Siglec-15–Fc | R&D Systems | 9227-SL-050 |
| AffiniPure Donkey anti-human IgG | Jackson ImmunoResearch | 709-605-098 |
| AffiniPure Goat anti-mouse IgG | Jackson ImmunoResearch | 115-605-071 |
| anti-GD2 (clone 14G2a) | BioLegend | 357302 or 357305 |
| CD163- | BioLegend | 155309 |
| CD11c- | BioLegend | 117337 |
| CD86- | BD | 563055 |
| CD11b | BioLegend | 101239 |
| I-A/I-E | BioLegend | 107643 |
| Ly6C | BioLegend | 128041 |
| CD45 | BioLegend | 103165 |
| PD-L1 | BioLegend | 124333 |
| CD80 | ThermoFisher | 46-0801-82 |
| Ly6G | ThermoFisher | 25-5931-82 |
| CD68 | BioLegend | 137003 |
| CD206 | BioLegend | 141733 |
| F4/80 | BioLegend | 123117 |

### Macrophage generation and stimulation

Primary human donor-derived macrophages were generated as previously described^34^. Briefly, leukocyte reduction system (LRS) chambers from anonymous donors were obtained from the Stanford Blood Center. Peripheral monocytes were purified through successive density gradients using Ficoll (Sigma Aldrich) and Percoll (GE Healthcare). Monocytes were then differentiated into macrophages by 7-9 days of culture in IMDM + 10% AB human serum (Life Technologies).

### In vitro phagocytosis assay

For all in vitro phagocytosis assays tumor cells and human macrophages were co-cultured at a ratio of 2:1 in ultra-low-attachment 96-well U-bottom plates (Corning) in serum-free RMPI (Thermo Fisher Scientific). In brief, tumor cells had either endogenous fluorescence (lentivirally expressed GFP) or were labeled with CFSE (Invitrogen) by suspending cells in PBS (5µM working solution) as per manufacturer instructions for 20 minutes at 37°C protected from light and washed twice with 20 mL FBS containing media before co-culture. Anti-GD2 (dinutuximab, acquired from United Therapeutics, or anti-B7-H3 (enoblituzumab, acquired from MacroGenics Inc.) was added alone or in combination with anti-CD47 (clone B6H12, BioXcell) at a concentration of 10 μg/mL. Tumor cells and antibodies were incubated for 30 minutes in a humidified, 5% CO2 incubator at 37°C. The plate was washed two times and human macrophages were added to the plate. After a 1-2 hour incubation at 37°C, phagocytosis was stopped by washing with 4°C PBS and centrifugation at 1200 RPMI before the cells were stained with APC-labeled anti-CD11b (Clone M1/70, Biolegend) to identify human macrophages and Live/Dead stain (Fixable Viability Dye, eBioscience) to exclude dead cells. Assays were analyzed by flow cytometry on an LRSFortessa Analyzer (BD Biosciences). Phagocytosis was measured as the number of CD11b+, CFSE/GFP+ macrophages, quantified as a percentage of the total CD11b+ macrophages.

### Molecular Cloning, protein expression and purification of murine anti-CD47 (ALX301)

CV1, is a high affinity human SIRPa variant, that has been reported to bind mouse CD47 with affinity of 30 nM compared with wildtype SIRPa which does not bind mouse CD47^54^. To design a version of CV1 with further improved binding to mouse CD47, structural models of mouse CD47 and a high affinity SIRPa, CV1 were built based on reported FD6 complex structure (PDB: 4KJY). From the modeled structures, various amino acid residues were identified and predicted to be critical for binding to mouse CD47. These mutations were explored experimentally and ALX301 was generated using site directed mutagenesis (QuickChange Lightning, Stratagene) to introduce V6I, A27I, I31R, K53R, E54D, H56P, L66T, S98I to the SIRP*α* domain which was then subcloned and fused to mouse IgG1 N297A via GeneArt Seamless Plus Cloning Assembly Kit (Invitrogen) according to manufacturer’s instructions. The construct was expressed in 5L scale Expi293 cells (Invitrogen) in shake flasks. Expression cultures were grown five days at 37°C in 8% CO2 while shaking at 125 RPM. Supernatants were harvested via centrifugation and sterile filtered. The protein was affinity purified using an Akta Avant150 (GE Healthcare), bound to MabSelectSure LX resin (GE Healthcare), washed with 1x phosphate-buffered saline (1xPBS), eluted with 0.1Mcitric acid pH 3.3, neutralized with 10% v/v 1M sodium phosphate pH 8, dialyzed into 1x PBS followed by a gel filtration polishing step where the main peak was pooled and collected.

### Surface Plasmon Resonance (SPR)

All kinetics were determined on a Proteon XPR 36 instrument (Bio-Rad, Hercules, CA) at 25C. The running buffer was PBS pH 7.4 supplemented with 0.01% Tween-20. ALX301 was coupled to one ligand channel of a GLC sensor chip using standard amine coupling chemistry. Mouse CD47, expressed and purified in-house, was injected as analyte in a 5-membered, 3-fold dilution series with 100 nM top concentration using the “single-shot kinetics” method. The sixth analyte channel was used as a buffer blank. The flow rate was 100 uL/min, the association time was 1 min and the dissociation time was 10 min. The data were double-referenced using interspots and the buffer blank before fitting to a 1:1 Langmuir binding model with the Proteon Manager software (Version 3.1.0.6).

### Cloning, expression, and purification of dinutuximab and dinutuximab LALA-PG

The gWIZ vector with a BM40 signal peptide was used for protein expression. DNA encoding dinutuximab’s human IgG1 heavy chain with and without LALA-PG mutation^55^, along with DNA encoding dinutuximab’s light chain, were ordered from Integrated DNA Technologies (Coralville, IA). The two heavy chains and light chain were individually cloned into AscI/BamHI digested gWIZ vector using Gibson Assembly.

Plasmids were transfected into Expi293F cells (ThermoFisher Scientific, Waltham, MA) in a 1:1 ratio of heavy chain:light chain using Expifectamine according to manufacturer’s instructions. Five days post-transfection, supernatant was harvested, adjusted to pH 8.0, and sterile filtered. Dinutuximab and dinutuximab LALA-PG were then purified using recombinant protein A-Sepharose 4B (ThermoFisher Scientific, Waltham, MA), buffer exchanged into antibody buffer (20mM L-Histidine, 250mM Sucrose, 0.05% Polysorbate 20, pH 6.0) using Pierce Slide-A-Lyzer G2 Dialysis Cassettes (ThermoFisher Scientific, Waltham, MA), and concentrated using Amicon Centrifugal Filters (Millipore Sigma, Burlington, MA).

### 14G2aproduction

Anti-GD2 monoclonal antibody (14G2a) was produced using a hybridoma gifted by Dr. Paul Sondel (University of Wisconsin). Purified antibody was generated by BioXCell, following 0.2um filtration and protein A purification. The lot of antibody tested negative for murine pathogens and had an endotoxin level of <2EU/mg.

### Exogenous ganglioside coating

GD2 (Matreya) was stored as a 1 mM stock solution in PBS at -20 °C. Cells were washed once with FACS buffer and the concentration adjusted to 4×10^6^ cells/mL. Where indicated, cells were treated with a sialidase cocktail, comprising 100 nM each of *Vibrio cholerae* sialidase, *Arthrobacter ureafaciens* sialidase, and *Clostridium perfringens* sialidase, for 30 minutes at 37 °C. The sialidases were expressed and purified in-house as previously described^56^. Cells were washed once with FACS buffer, then twice with PBS to remove BSA, as the presence of any residual protein was found to decrease the efficiency of labeling. The cell concentration was adjusted to 4×10^6^ cells/mL in PBS, and stock GD2 was added to the cell suspension to achieve a final concentration of 200 μM.

The solution was mixed by pipetting and incubated for 30 minutes on ice. The cells were washed twice with FACS buffer prior to subsequent analyses.

### Siglec ligand detection

Siglec ligands were quantified via flow cytometry. 4 μg/mL of each Siglec-Fc (R&D Systems) was pre-complexed with either 8 μg/mL AffiniPure Donkey anti-human IgG Alexa Fluor 647 (Jackson ImmunoResearch) or AffiniPure Goat anti-mouse IgG Alexa Fluor 647 (Jackson ImmunoResearch) in FACS buffer (0.5% bovine serum albumin in phosphate buffered saline) for 30 minutes on ice. Cells were washed once with FACS buffer then resuspended at 4×10^6^ cells/mL. Where indicated, GD2 was blocked by staining with 10 ug/mL anti-GD2 antibody (clone 14G2a; BioLegend) for 30 minutes on ice. Cells were again washed in FACS buffer and resuspended in 100 μL of the pre-complexed staining solution at 4×10^6^ cells/mL and incubated for 30 minutes on ice. The cells were then washed twice with FACS buffer. Dead cells were labeled using SytoxBlue (Thermo Fisher Scientific) prior to performing flow cytometry on an LSR II (BD Biosciences).

### In vivo experiments

#### Mice

All animal studies were carried out under protocols approved Stanford University and University of California San Francisco Animal Care and Use Committees. Immunodeficient NSG (NOD.Cg-Prkdcscid Il2rgtm1Wjl/SzJ) or immunocompetent 129SvJ mice were ordered from JAX or bred in house. TH-MYCN mice were bred in house as previously described^22^. Mice used for *in vivo* experiments were between 6 and 12 weeks old.

### Neuroblastoma *in vivo* models

CHLA255 or KCNR tumor cells expressing green fluorescence protein and luciferase were expanded under standard culture conditions (described above) and harvested with Trypsin (Gibco, Thermo Fisher Scientific). For the orthotopic experiments 1e6 KCNR-GFP Luc or 1e6 CHLA255-GFP Luc tumor cells were implanted into the renal capsule as previously descibed^57^. Tumor growth was followed by bioluminescence intensity (BLI) on an IVIS Spectrum In Vivo Imaging System (PerkinElmer) 4 minutes after 3 mg d-luciferin (PerkinElmer) was injected intraperitoneally. BLI values were quantified with the Live Image v.4.0 software (Living Image; PerkinElmer). Four days after tumor implantation, mice were randomized based on BLI and antibody treatment was initiated. 400µg IgG control (BioXcell), 400µg anti-CD47 (clone B6.H12, BioXcell) or 400µg magrolimab (Hu5F9-G4, Forty Seven, Inc./Gilead), 300µg anti-B7-H3 (Enoblituzumab, MacroGenics Inc.), 300µg anti-GD2 (Dinutuximab, United Therapeutics), or combination treatments were administered intraperitoneally every other day for three doses. For the metastatic model experiment 1e6 CHLA255-GFP Luc tumor cells were injected into the tail vein and randomization and antibody treatment was performed as in the orthotopic models. For the Fc-dead experiment, 400µg IgG control (BioXcell), 400µg anti-CD47 (clone B6.H12, BioXcell), 300µg Fc-dead anti-GD2 (Dinutuximab LALA-PG) or 300µg FC-active anti-GD2 (Dinutuximab) (generated as described above), or combination treatments were administered intraperitoneally twice in three days. Mice received two doses compared to three doses of the other antibody experiments performed in that model because of limited availability of the amount of Fc-dead anti-GD2.

### Osteosarcoma *in vivo* models

MG63.3 osteosarcoma cells were expanded under standard culture conditions (described above) and harvested with Trypsin (Gibco, Thermo Fisher Scientific). For the orthotopic model, 1e6 MG63.3 cells were injected periosteal to the tibia of NSG mice. Antibody treatment with 400µg IgG control (BioXcell), 400µg anti-CD47 (clone B6.H12, BioXcell), 300µg anti-B7-H3 (enoblituzumab, MacroGenics Inc.), 300 µg anti-GD2 (dinutuximab, United Therapeutics), or combination treatments were administered intraperitoneally three times a week starting seven days after tumor engraftment for four weeks. Tumor growth was measured with digital calipers once to twice weekly, and the tumor area was calculated by multiplying the lengths of the major and minor axes. Mice were euthanized when tumors exceeded a size set by institutional protocol.

For the metastatic model, orthotopic tumors were implanted as described above and allowed to grow for thirty days. The tumor-bearing leg of all mice was amputated using sterile technique under isoflurane anesthesia. Buprenorphine 0.05 mg/kg was injected subcutaneously for pain control. Mice were randomized to the treatment groups described above, based on their pre-amputation tumor sizes (groups were statistically identical). Antibody treatment was provided for a total course of 4 weeks. Thirty days after amputation, experimental mice were euthanized and lungs were removed for evaluation of metastases. Lung metastasis were quantified by counting the number of visible GFP+ lesions on an ultraviolet gel box. Tissues were then formalin-fixed and further analyzed by IHC.

### H69 Human SCLC in vivo Model

NCI-H69 SCLC cells were collected, counted, and resuspended in media without antibiotics in single tumor aliquots of 2e6 cells in 100 ul to ensure that each tumor contained the identical number of cells. Cells were mixed with Matrigel (Fisher Scientific) at 1:1 ratio and then injected into both hind flanks of NSG mice. When tumors were palpable mice were randomized and treated with an isotype control, anti-CD47, ant-GD2, or both anti-CD47 and anti-GD2 as described for the models above. Tumors were measured with digital calipers twice weekly and mice were euthanized when tumors reached the volume threshold set by institutional protocol.

### Syngeneic TH-MYCN in vivo Model

De novo tumors from TH-MYCN mice were harvested, dissociated into single cells, and 1e6 cells were injected subcutaneously in the right flank of female 129SvJ mice (8 weeks old). The cells were resuspended in a 1:1 solution of Neurobasal media (Gibco) and Geltrex (Gibco) prior to injection. After tumor formation, tumor size was measured twice a week. Mice were enrolled for treatment when tumors reached a volume of ∼700 mm^3^. Mice were treated by intraperitoneal injection with 50ug of murine anti-GD2 antibody (14g2a, see details above), 30mg/kg of murine CD47 blocker (ALX301) or a combination of both twice a week, for a total of three weeks. Mice were sacrificed when the tumor size reached 2cm in one direction. For macrophage depletion, mice were pre-treated by intravenous injection with 100uL of clodronate liposomes (Liposoma), followed by 400ug of anti-CSF1R (BioXcell, AFS98) by intraperitoneal injection two days later, then enrolled in treatment groups. Mice were continued to be treated with 400ug of anti-CSF1R three times a week for the three weeks of treatment.

### Neutrophil Depletion

The metastatic CHLA255 model was used as described above. 500ug anti-Ly6G (clone 1A8, BioXcell) was injected intaperitoneally every other day throughout the course of treatment, beginning two days prior to the start of treatment. To confirm depletion of neutrophils, peripheral blood was collected on the final day of treatment, ACK lysed, and stained with anti-Ly6G/Ly6C (clone RB6-8C5, ThermoFisher).

### Phenotyping of intratumoral macrophages

1e6 MG63.3 tumor cells were injected as described above and allowed to grow for three weeks. Antibody treatment was administered intraperitoneally with 400ug IgG control, 300ug anti-GD2 (Dinutuximab) and/or 400ug anti-CD47 (B6H12) three times per week for one week. Tumor tissues were excised and split into half (other half used for immunohistochemistry, see below). Tumors were homogenized into single cells in complete media with a Miltenyi GentleMACS. Digested tumors were subsequently passed through a 70 μM filter and adjusted to a concentration of 10e6 cells/mL for staining in FACS buffer. Dead cells were stained with Live/Dead Violet (ThermoFisher) in 1 mL PBS. Cells were subsequently blocked with 2 μL of TruStain FcX Fc receptor blocking antibody (BioLegend) for 10 minutes on ice. Cell suspensions were washed and resuspended in FACS buffer (PBS + 5% FBS + 5mM EDTA) with the following fluorophore-labelled antibodies for 30 minutes on ice: CD163-BV421, CD11c-BV510, BV605-CD86, CD11b-BV650, I-A/I-E-BV711, Ly6C-BV785, CD45-Spark Blue 550, PD-L1-PerCP-Cy5.5, CD80, PerCP-ef710, Ly6G-PE-Cy7, CD68-AF647, CD206-AF700, F4/80-APC-Cy7. The proportions of M2-like TAMs (CD45^+^CD11b^+^F4/80^+^CD163^+^CD206^+^) within the tumor tissues of mice receiving different treatments were analyzed. The same tumors were also sent

### Immunohistochemistry (IHC) and analysis

For IHC analysis of orthotopic xenograft osteosarcoma tumors, tumors were obtained as above. Tumors were harvested for further analysis for F4/80 and iNOS positivity. For visualization of lung metastasis, lung tissue was harvested from mice treated as described under the metastatic osteosarcoma model and GFP+ tumor cells were detected via GFP staining. Formalin-fixed, paraffin-embedded xenograft tumor sections were used. GFP (Abcam, ab6556, 1:1500) and F4/80 (Cell Signalling, D2S9R) staining was performed manually and iNOS (Novus NB300-605SS, 1:500 dilution) staining was performed using the Ventana Discovery platform. In brief, tissue sections were incubated in either 6mM citrate buffer (GFP and F4/80) or Tris EDTA buffer (iNOS) (cell conditioning 1; CC1 standard) at 100 degrees for 25minutes (GFP/F480) or 95°C for 1 hour (iNOS) to retrieve antigenicity, followed by incubation with the respective primary antibody for one hour. Bound primary antibodies were incubated with the respective secondary antibodies (Vector Laboratories or Jackson Laboratories) with 1:500 dilution, followed by Ultramap HRP and Vectore Lab (GFP/F480) or Chromomap DAB (iNOS) detection. For IHC analysis, lung macrometastases (> 200 um) were identified based on GFP positivity. F4/80 and iNOS positivity was analyzed for each tumor (each area ∼ 3 mm^2^). F4/80 and iNOS IHC positivity scores were automatically quantified in the regions of interest with Aperio ImageScope software. Regions of interest were randomly selected within the tumor to exclude macrophages present in the normal tissue around the tumor.

### Magnetic resonance imaging (MRI)

MRI was performed to investigate the effect of anti-GD2 and anti-CD47 in stimulating macrophage infiltration. 1e6 MG63.3 tumor cells were injected as described above and allowed to grow for three weeks. At this time mice underwent a baseline MRI. A second MRI was performed after antibody treatment with 300ug anti-GD2 (Dinutuximab) and 400ug anti-CD47 (B6H12) or 400ug IgG control three times per week for one week. On the same day after the second time MRI, the mice received ferumoxytol intravenously at the dose of 0.5 mmol Fe/kg. Repeat MRI was performed 12 hours after ferumoxytol injection. MRI studies were performed on a 7T MR scanner (Bruker Biospin, Billerica, MA), using a 2.9-cm-inner-diameter (4 cm active length), Millipede RF coil (ExtendMR LLC, Milpitas, CA) operating in quadrature and tuned to 298.06 MHz, specifically designed for high field micro imaging of the mouse, and the following pulse sequences with a field of view of 3 cm x 3 cm and a slice thickness of 0.75 mm: T2-weighted fast spin echo (FSE): repetition time (TR): 2528 ms, echo time (TE): 33 ms and T2*-map multi-gradient echo (MGE): Flip angle: 80°, TR: 999 ms, TE: 3, 7,12,16, 20, 25, 29, 33, 38 and 42 ms. Mean T2* values of the tumors were analyzed using the Osirix software version 8.0.2 (Pixmeo SARL, Bernex, Switzerland) to quantify tumor contrast enhancement. The slice with the largest cross-sectional area of the tumor in the T2-weighted FSE image volume was identified for each animal. The visible tumor boundary was defined within that slice. The mean T2* value within the visible tumor boundary was calculated for each animal. T2* means were averaged across control and treated animal groups at each timepoint.

### Cell death and Calreticulin assessment by Flow cytometry

100,000 neuroblastoma tumor cells were plated in triplicate in a 96 well plate (Corning) and treated with 10ug/ml anti-GD2 (Dinutuximab, acquired from United Therapeutics) for indicated timepoints at 37°C. Cells were washed twice before staining with anti-calreticulin (Abcam) for 30 minutes on ice protected from light. Afterwards, cells were washed with FACS buffer and stained with DAPI, before data collected with an LSR Fortessa X-20.

### Statistical analysis

Data analysis and visualization was performed using Prism 8.4 (GraphPad Software). Graphs represent either group mean values ± s.d. (for *in vitro* experiments) or ± s.e.m. (for in vivo experiments) or individual values. For in vitro studies, statistical comparisons were made with either unpaired t-tests when comparing two groups or a one-way ANOVA with multiple comparison correction when comparing more than two groups. For in vivo studies, survival curves were compared with the log-rank test, tumor growth was compared with repeated-measures ANOVA, and the Mann-Whitney-test was used to compare two groups. *P* < 0.05 was considered statistically significant. *P* values are denoted with asterisks: *P* > 0.05, NS; \**P* < 0.05; \*\**P* < 0.01; \*\*\**P* < 0.001; and \*\*\*\**P* < 0.0001.

## Acknowledgements

This work was supported by an Alex’s Lemonade Stand ‘A’ Award (RGM) and NIH P01 CA217959 (CLM, WAW, RGM). RGM is the Taube Distinguished Scholar for Pediatric Immunotherapy at Stanford University School of Medicine. JT is supported by German Cancer Aid (Deutsche Krebshilfe) grant no. P-91650709. This work was supported by the National Cancer Institute (R01-CA227942 to CRB and F30-CA232541 to BAHS and U01-CA217864 to WAW, U01-CA213273 and R35-CA231997 to JS), the American Cancer Society (post-doctoral fellowship to GLC). B.A.H.S is supported by the Stanford School of Medicine Medical Scientist Training Program (T32-GM007365). M.H.L is supported by a Blavatnik Family Fellowship. P.L.L. is supported by the Stanford Bio-X Bowes Fellowship and the Stanford Medical Scientist Training Program. We thank the Stanford Neuropathology Department for their help with immunohistochemistry. All cartoons were created with BioRender.com.

---

**Extended Data Figure 1.**
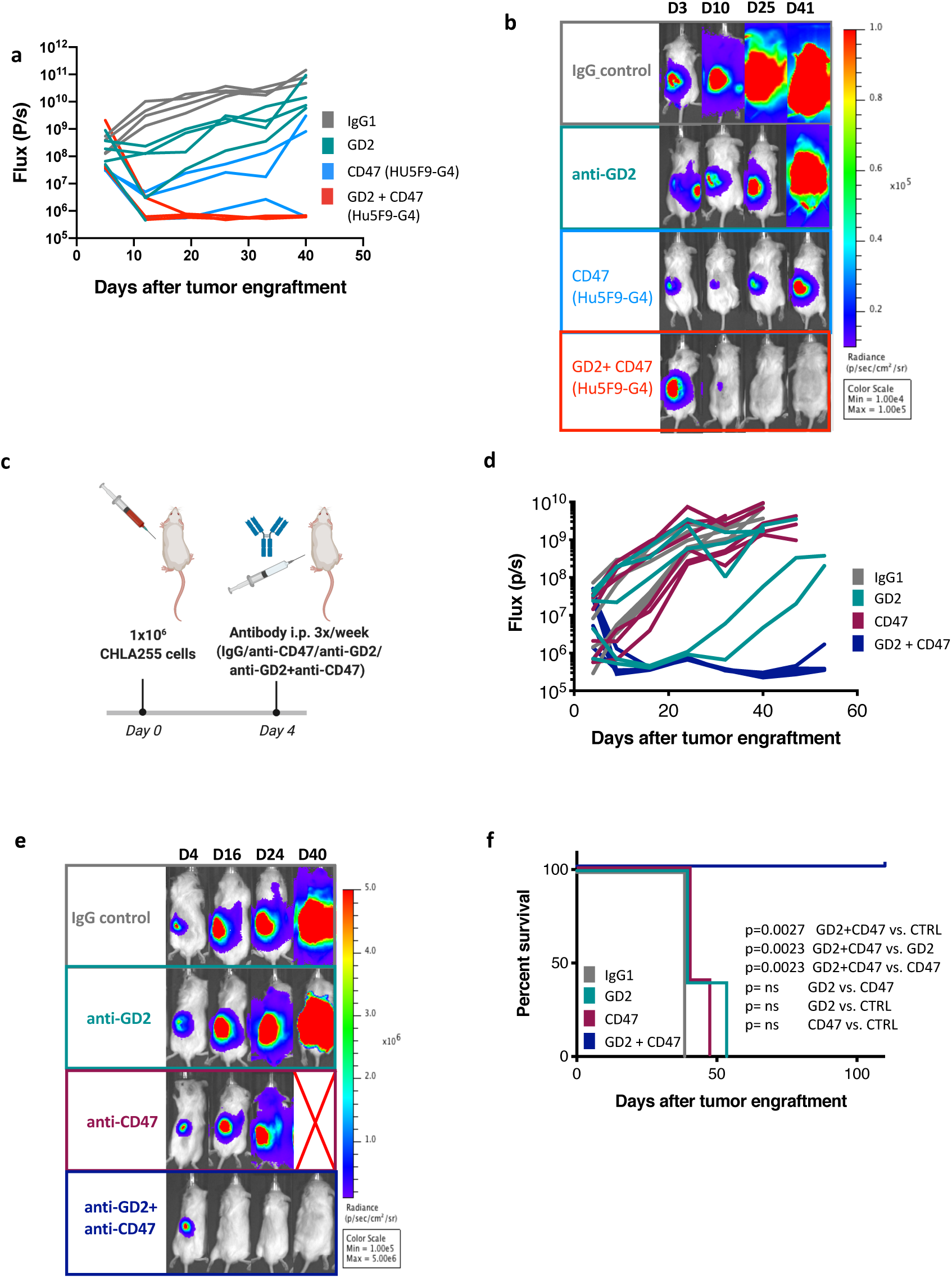
Anti-CD47 and anti-GD2 synergize to mediate significant anti-tumor activity in orthotopic models of neuroblastoma. One million KCNR neuroblastoma cells expressing GFP-luciferase were implanted into the renal capsule and treated four days later with IgG control, anti-G2 mAb, anti-CD47 (magrolimab, Hu5F9-G4) mAb or dual anti-GD2/anti-CD47 (Hu5F9-G4) every other day for three doses as in Figure 1c. **a**, Quantification of tumor progression for each individual mouse as measured by flux values acquired via bioluminescence (BLI) photometry. **b**, BLI images of representative mice from each treatment group at different time points. Experiment was performed one time. N=5 mice per group for IgG and anti-GD2 and those groups are the same as in Figure 1c-f. N=3 for anti-CD47 and anti-CD47 + anti-GD2. **c**, One million MYCN non-amplified CHLA255 neuroblastoma cells expressing GFP-luciferase cells were implanted into the renal capsule in NSG mice. Four days later, mice were injected with IgG control, anti-GD2 mAb, anti-CD47 mAb or dual anti-GD2/anti-CD47 treatment every other day for three doses. d, Quantification of tumor progression for each individual mouse as measured by flux values acquired via BLI photometry. **e,** BLI images of representative mice from each treatment group shown in d at different time points. Red cross indicates deceased mouse. **f,** Survival curves for mice bearing tumors shown in **c**. Representative data from three independent experiments. N=5 mice per group. Survival curves were compared using the log-rank test.

**Extended Data Figure 2.**
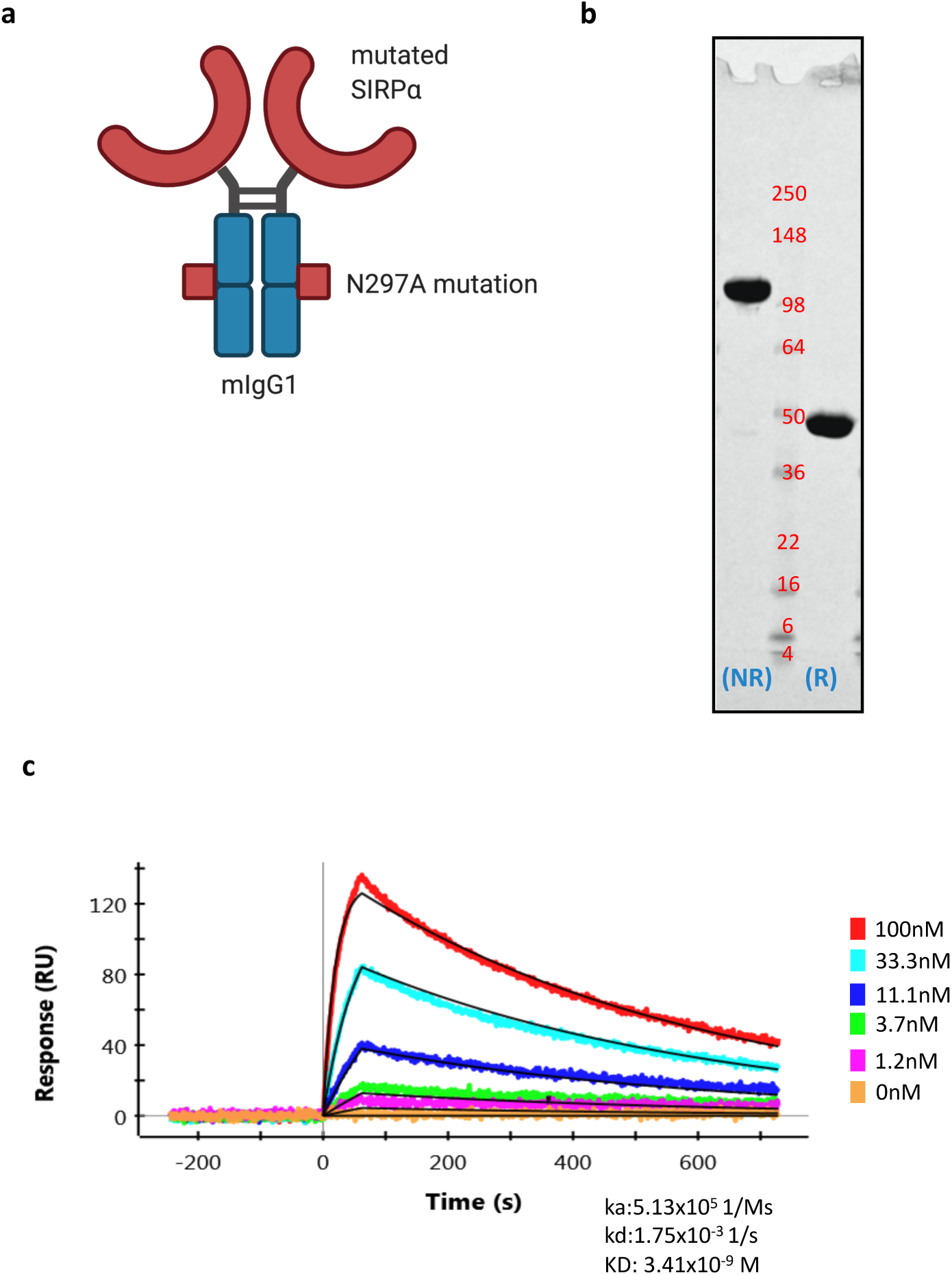
Expression and affinity of ALX301, a novel molecule capable of blocking murine CD47. **a,** Schematic of ALX301: Murine IgG1 containing a N297A mutation was fused to a mutated SIRP*α* capable of binding murine CD47 with enhanced affinity. **b,** Purified ALX301 was detected on 4-20% Tris-gylcine gel in non-reducing (NR) and reducing (R) buffer. ALX301 runs slightly larger than the expected 76.16 kDa (NR) and 38.038 kDa (R). **c,** ALX301 was immobilized on GLC sensor chip (Bio-rad). Recombinant mouse CD47 protein was injected as analyte over the chip at 5 concentrations, 3-fold dilution (100nM, 33.3nM, 11.1nM, 3.7nM, 1.2nM). Using Langmuir binding model for curve fitting, the binding of ALX301 to mouse SIRPa was determined to be 3.41nM. Association rate (ka), dissociation rate (kd), affinity (KD).

**Extended Data Figure 3.**
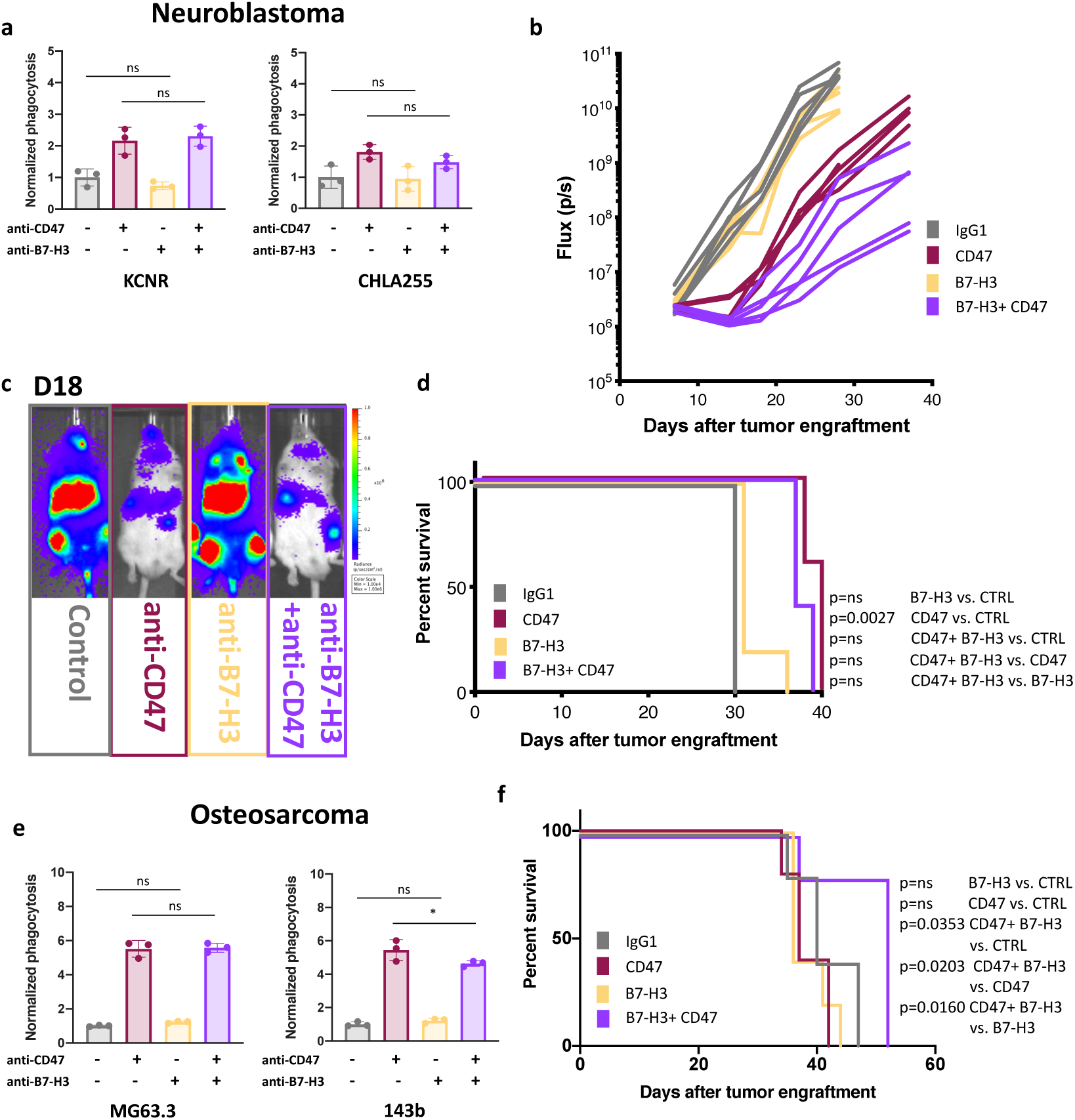
Anti-B7-H3 does not synergize with anti-CD47 in xenograft models of neuroblastoma or osteosarcoma. **a,** Graphs show flow cytometry-based quantification of phagocytosis of KCNR (left) and CHLA255 (right) neuroblastoma cell lines co-cultured with human blood derived macrophages in the presence of anti-CD47 mAb, anti-B7-H3 mAb or dual treatment, compared with untreated control; results normalized to the phagocytosis in the untreated control in each independent replicate experiment, for each cell line and blood donor. Error bars, s.d. of three experimental replicates. Statistical comparisons performed with one-way ANOVA with multiple comparisons correction, *P < 0.05, **P < 0.01, ***P < 0.001, \*\*\*\**P* < 0.0001, ns P>0.05. Representative data from at least three experiments with three different blood donors. **b,** One million CHLA255 neuroblastoma cells expressing GFP-luciferase cells were engrafted into NSG mice by tail-vein injection. Four days later, mice were injected with IgG control, anti-CD47 mAb, anti-B7-H3 mAb, or dual treatments every other day for three doses. Quantification of tumor progression for each individual mouse as measured by flux values acquired via bioluminescence (BLI) photometry. **c,** BLI images of representative mice from each treatment group shown in **b** at one time point. **d,** Survival curves for mice bearing tumors shown in **b.** Survival curves were compared using the log-rank test. N=5 mice per group. **e,** Graph shows flow cytometry-based quantification of phagocytosis of MG63.3 (left) and 143b (right) osteosarcoma cell lines co-cultured with human blood derived macrophages in the presence of anti-CD47 mAb, anti-B7-H3 mAb or dual treatment, compared with untreated control; results normalized to the phagocytosis in the untreated control in each independent replicate experiment, for each cell line and blood donor. Error bars, s.d. of three experimental replicates. Statistical comparisons performed as in **a**. Representative data from at least three experiments with three different blood donors. **f,** Survival curves for mice that received hind leg injection of one million MG63.3 cells. Seven days later, mice were injected with IgG control, anti-CD47 mAb, anti-B7-H3 mAb or dual treatments 3 times/week for 4 weeks. Survival curves were compared using the log-rank test. N=5 mice per group.

**Extended Data Figure 4.**
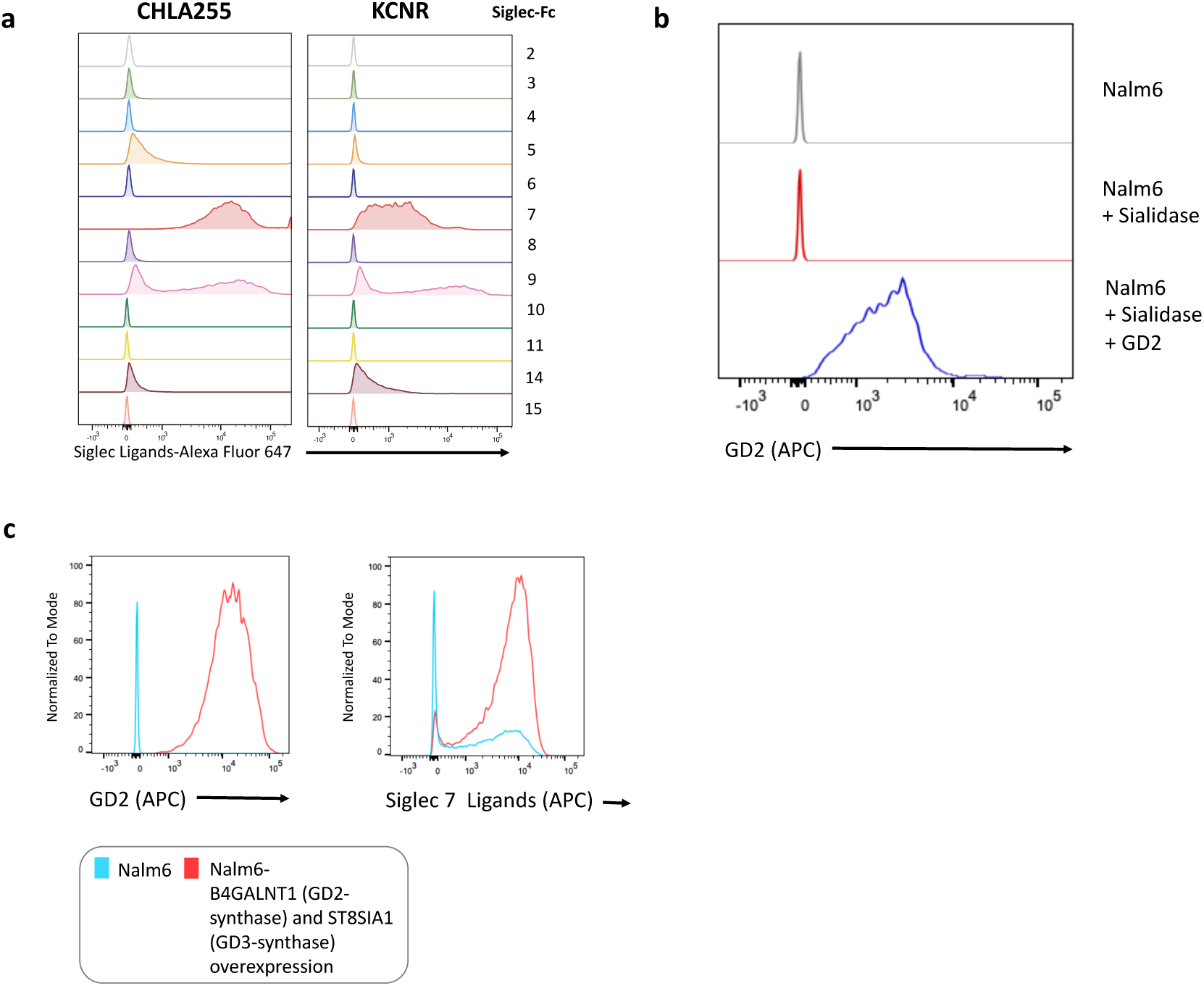
Siglec binding on neuroblastoma cells and coating/forced expression of GD2 on GD2 negative Nalm6. **a,** Flow cytometric histograms of CHLA255 and KCNR stained with soluble versions of human Siglecs. **b,** Flow cytometric histograms showing intensity of GD2 staining of Nalm6 cells (top), Nalm6 cells treated with sialidases (middle) and Nalm6 cells treated with sialidases and coated with GD2 (bottom) **c,** Flow cytometric analysis of the expression of GD2 (left panel) and Siglec-7 ligands (right panel) on the surface of wildtype Nalm6 cells or Nalm6 cells transduced with lentiviral constructs expressing B4GALNT1 (GD2-synthase) and ST8SIA1 (GD3-synthase).

**Extended Data Figure 5.**
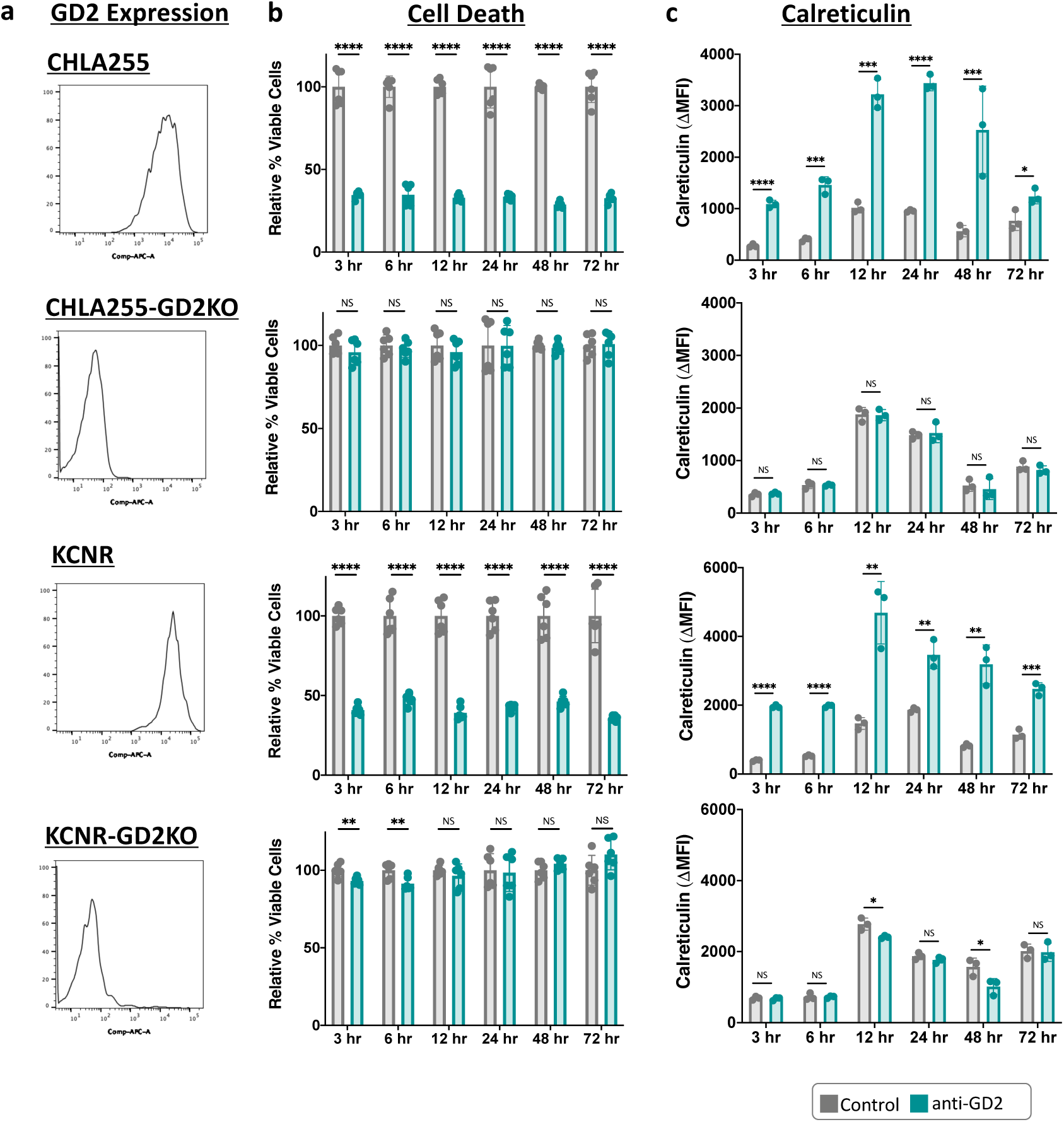
Upregulation of ‘Eat me’ signal calreticulin and induction of cell death by ligation with anti-GD2 mAb on neuroblastoma cell lines. **a,** Flow cytometric analysis of the levels of expression of GD2 on the surface of neuroblastoma cell lines CHLA255 and KCNR and their isogenic GD2-KO (B4GALNT1 KO) versions. **b,** Flow-based quantification of cell viability. Neuroblastoma cell lines and their isogenic GD2-KO versions were incubated with anti-GD2 mAb for 12 hours at 37 degrees and stained with DAPI. Percent of DAPI-populations were normalized to the untreated control for each cell line. Error bars, s.d. of 6 experimental replicates. Statistical comparisons performed with the unpaired t-test, *P < 0.05, **P < 0.01, ***P < 0.001, \*\*\*\**P* < 0.0001, ns P>0.05. **c,** Graph shows flow cytometric quantification of the expression of calreticulin (ΔMFI) on the surface of neuroblastoma cell lines treated as in b. Error bars, s.d. of *n* = 3 experimental replicates. ΔMFI was calculated as the difference between the MFI in the PE channel of the calreticulin stained sample and the MFI in the PE channel of an isotype stained sample from the same experimental condition. Error bars, s.d. of 3 experimental replicates. Statistical comparisons performed as in **b.** The full timecourse experiment was performed once and the twelve-hour timepoint is identical to figure 4h.

**Extended Data Figure 6.**
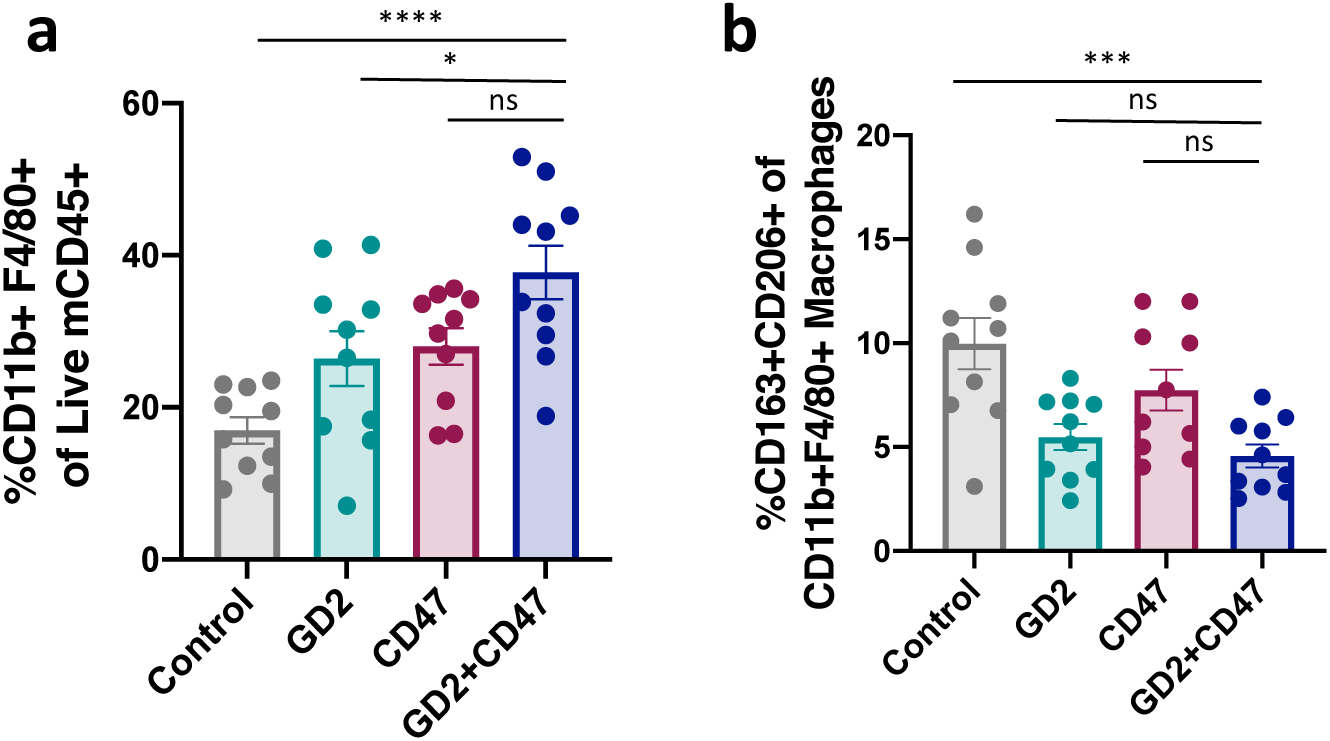
Anti-GD2 and anti-CD47 treatment leads to increase in macrophage infiltration and reduction of M2 macrophages. **a-b,** Flow cytometric analysis of a, overall macrophage infiltration or **b,** M2 macrophages (defined by CD163+ and CD206+, gated on CD45+, CD11b+, F4/80 + macrophages) in tumors harvested after seven days of treatment with indicated mAb, pooled results from two independent experiments (n=10 per group). Error bars, s.e.m. of ten biologic replicates. Statistical comparisons were made between groups with the one-way ANOVA with correction for multiple comparisons correction, *P < 0.05, **P < 0.01, ***P < 0.001, \*\*\*\**P* < 0.0001, ns P>0.05.

**Extended Data Figure 7.**
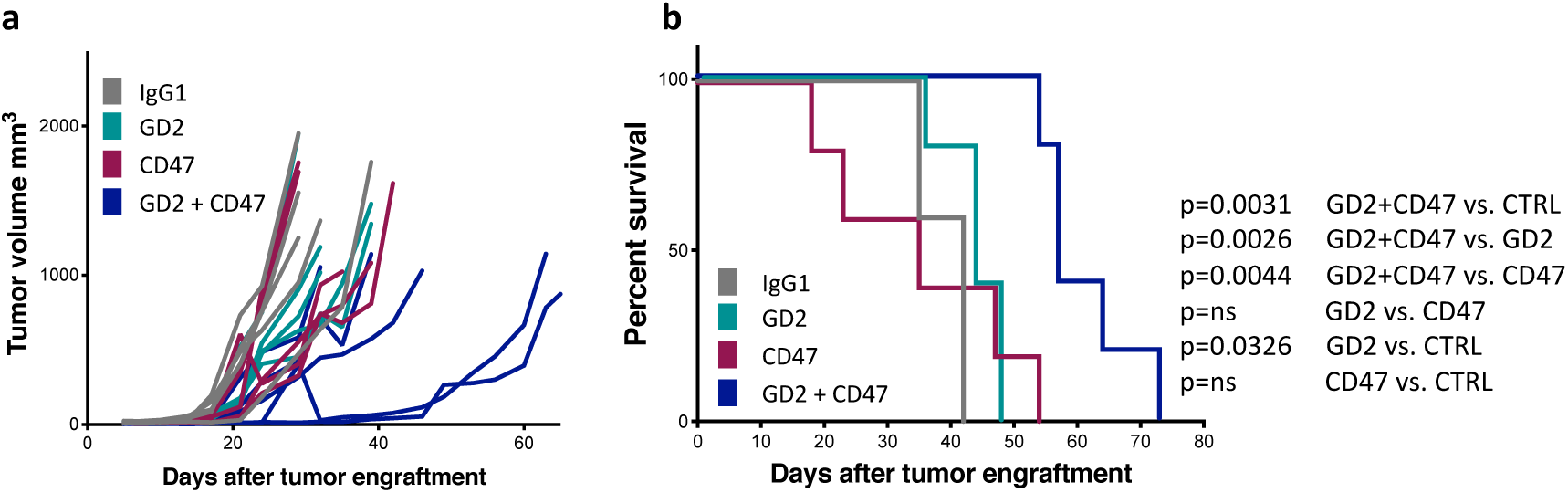
Anti-CD47 and anti-GD2 synergize to mediate significant anti-tumor activity in xenograft model of small cell lung cancer (SCLC). **a,** NCI-H69 SCLC were engrafted on both flanks of NSG mice. Quantification of tumor growth for each individual tumor was assessed by caliper measurement. **b,** Survival curves for mice bearing tumors shown in **a.** Survival curves were compared using the log-rank test. Representative of two independent experiments.

## Data availability

The datasets generated during this study are available from the corresponding author upon reasonable request.

## References

1. Saito, M., Yu, R. K. & Cheung, N. K. Ganglioside GD2 specificity of monoclonal antibodies to human neuroblastoma cell. Biochem Biophys Res Commun 127, 1–7, doi:10.1016/s0006-291x(85)80117-0 (1985).

2. Schulz, G. et al. Detection of ganglioside GD2 in tumor tissues and sera of neuroblastoma patients. Cancer Res 44, 5914–5920 (1984).

3. Long, A. H. et al. Reduction of MDSCs with All-trans Retinoic Acid Improves CAR Therapy Efficacy for Sarcomas. Cancer Immunol Res 4, 869–880, doi:10.1158/2326-6066.CIR-15-0230 (2016).

4. Roth, M. et al. Ganglioside GD2 as a therapeutic target for antibody-mediated therapy in patients with osteosarcoma. Cancer 120, 548–554, doi:10.1002/cncr.28461 (2014).

5. Dobrenkov, K., Ostrovnaya, I., Gu, J., Cheung, I. Y. & Cheung, N. K. Oncotargets GD2 and GD3 are highly expressed in sarcomas of children, adolescents, and young adults. Pediatr Blood Cancer 63, 1780–1785, doi:10.1002/pbc.26097 (2016).

6. Mount, C. W. et al. Potent antitumor efficacy of anti-GD2 CAR T cells in H3-K27M(+) diffuse midline gliomas. Nat Med 24, 572–579, doi:10.1038/s41591-018-0006-x (2018).

7. Cheresh, D. A., Rosenberg, J., Mujoo, K., Hirschowitz, L. & Reisfeld, R. A. Biosynthesis and expression of the disialoganglioside GD2, a relevant target antigen on small cell lung carcinoma for monoclonal antibody-mediated cytolysis. Cancer Res 46, 5112–5118 (1986).

8. Battula, V. L. et al. Ganglioside GD2 identifies breast cancer stem cells and promotes tumorigenesis. J Clin Invest 122, 2066–2078, doi:10.1172/JCI59735 (2012).

9. Yu, A. L. et al. Anti-GD2 antibody with GM-CSF, interleukin-2, and isotretinoin for neuroblastoma. N Engl J Med 363, 1324–1334, doi:10.1056/NEJMoa0911123 (2010).

10. Ladenstein, R. et al. Interleukin 2 with anti-GD2 antibody ch14.18/CHO (dinutuximab beta) in patients with high-risk neuroblastoma (HR-NBL1/SIOPEN): a multicentre, randomised, phase 3 trial. Lancet Oncol 19, 1617–1629, doi:10.1016/S1470-2045(18)30578-3 (2018).

11. Cheung, N. K. et al. Murine anti-GD2 monoclonal antibody 3F8 combined with granulocyte-macrophage colony-stimulating factor and 13-cis-retinoic acid in high-risk patients with stage 4 neuroblastoma in first remission. J Clin Oncol 30, 3264–3270, doi:10.1200/JCO.2011.41.3807 (2012).

12. Matthay, K. K., et al. Neuroblastoma. Nat Rev Dis Primers 2, 16078, doi:10.1038/nrdp.2016.78 (2016).

13. Hobbie, W. L. et al. Late effects in survivors of tandem peripheral blood stem cell transplant for high-risk neuroblastoma. Pediatr Blood Cancer 51, 679–683, doi:10.1002/pbc.21683 (2008).

14. Suh, E. et al. Late mortality and chronic health conditions in long-term survivors of early-adolescent and young adult cancers: a retrospective cohort analysis from the Childhood Cancer Survivor Study. Lancet Oncol 21, 421–435, doi:10.1016/S1470-2045(19)30800-9 (2020).

15. Armstrong, A. E. et al. Late Effects in Pediatric High-risk Neuroblastoma Survivors After Intensive Induction Chemotherapy Followed by Myeloablative Consolidation Chemotherapy and Triple Autologous Stem Cell Transplants. J Pediatr Hematol Oncol 40, 31–35, doi:10.1097/MPH.0000000000000848 (2018).

16. Hingorani, P. et al. Phase II study of antidisialoganglioside antibody, dinutuximab, in combination with GM-CSF in patients with recurrent osteosarcoma (AOST1421): A report from the Children’s Oncology Group. Journal of Clinical Oncology 38, 10508–10508, doi:10.1200/JCO.2020.38.15_suppl.10508 (2020).

17. Edelman, M. et al. The anti-disialoganglioside (GD2) antibody dinutuximab (D) for second-line treatment (2LT) of patients (pts) with relapsed/refractory small cell lung cancer (RR SCLC): Results from part II of the open-label, randomized, phase II/III distinct study. Journal of Clinical Oncology 38, 9017–9017, doi:10.1200/JCO.2020.38.15_suppl.9017 (2020).

18. Grant, S. C. et al. Targeting of small-cell lung cancer using the anti-GD2 ganglioside monoclonal antibody 3F8: a pilot trial. Eur J Nucl Med 23, 145–149, doi:10.1007/BF01731837 (1996).

19. Jaiswal, S. et al. CD47 is upregulated on circulating hematopoietic stem cells and leukemia cells to avoid phagocytosis. Cell 138, 271–285, doi:10.1016/j.cell.2009.05.046 (2009).

20. Sikic, B. I., et al. First-in-Human, First-in-Class Phase I Trial of the Anti-CD47 Antibody Hu5F9-G4 in Patients With Advanced Cancers. J Clin Oncol 37, 946–953, doi:10.1200/JCO.18.02018 (2019).

21. Advani, R. et al. CD47 Blockade by Hu5F9-G4 and Rituximab in Non-Hodgkin’s Lymphoma. N Engl J Med 379, 1711–1721, doi:10.1056/NEJMoa1807315 (2018).

22. Weiss, W. A., Aldape, K., Mohapatra, G., Feuerstein, B. G. & Bishop, J. M. Targeted expression of MYCN causes neuroblastoma in transgenic mice. EMBO J 16, 2985–2995, doi:10.1093/emboj/16.11.2985 (1997).

23. Sockolosky, J. T. et al. Durable antitumor responses to CD47 blockade require adaptive immune stimulation. Proc Natl Acad Sci U S A 113, E2646–2654, doi:10.1073/pnas.1604268113 (2016).

24. Willingham, S. B. et al. The CD47-signal regulatory protein alpha (SIRPa) interaction is a therapeutic target for human solid tumors. Proc Natl Acad Sci U S A 109, 6662–6667, doi:10.1073/pnas.1121623109 (2012).

25. Erbe, A. K. et al. Neuroblastoma Patients’ KIR and KIR-Ligand Genotypes Influence Clinical Outcome for Dinutuximab-based Immunotherapy: A Report from the Children’s Oncology Group. Clin Cancer Res 24, 189–196, doi:10.1158/1078-0432.CCR-17-1767 (2018).

26. Forlenza, C. J. et al. KIR3DL1 Allelic Polymorphism and HLA-B Epitopes Modulate Response to Anti-GD2 Monoclonal Antibody in Patients With Neuroblastoma. J Clin Oncol 34, 2443–2451, doi:10.1200/JCO.2015.64.9558 (2016).

27. Wei, J. S. et al. Clinically Relevant Cytotoxic Immune Cell Signatures and Clonal Expansion of T-Cell Receptors in High-Risk MYCN-Not-Amplified Human Neuroblastoma. Clin Cancer Res 24, 5673–5684, doi:10.1158/1078-0432.CCR-18-0599 (2018).

28. Asgharzadeh, S. et al. Clinical significance of tumor-associated inflammatory cells in metastatic neuroblastoma. J Clin Oncol 30, 3525–3532, doi:10.1200/JCO.2011.40.9169 (2012).

29. Song, L. et al. Valpha24-invariant NKT cells mediate antitumor activity via killing of tumor-associated macrophages. J Clin Invest 119, 1524–1536, doi:10.1172/JCI37869 (2009).

30. Loo, D. et al. Development of an Fc-enhanced anti-B7-H3 monoclonal antibody with potent antitumor activity. Clin Cancer Res 18, 3834–3845, doi:10.1158/1078-0432.CCR-12-0715 (2012).

31. Crocker, P. R., Paulson, J. C. & Varki, A. Siglecs and their roles in the immune system. Nat Rev Immunol 7, 255–266, doi:10.1038/nri2056 (2007).

32. Avril, T., Floyd, H., Lopez, F., Vivier, E. & Crocker, P. R. The membrane-proximal immunoreceptor tyrosine-based inhibitory motif is critical for the inhibitory signaling mediated by Siglecs-7 and -9, CD33-related Siglecs expressed on human monocytes and NK cells. J Immunol 173, 6841–6849, doi:10.4049/jimmunol.173.11.6841 (2004).

33. Yamaji, T., Mitsuki, M., Teranishi, T. & Hashimoto, Y. Characterization of inhibitory signaling motifs of the natural killer cell receptor Siglec-7: attenuated recruitment of phosphatases by the receptor is attributed to two amino acids in the motifs. Glycobiology 15, 667–676, doi:10.1093/glycob/cwi048 (2005).

34. Barkal, A. A. et al. CD24 signalling through macrophage Siglec-10 is a target for cancer immunotherapy. Nature 572, 392–396, doi:10.1038/s41586-019-1456-0 (2019).

35. Tsao, C. Y. et al. Anti-proliferative and pro-apoptotic activity of GD2 ganglioside-specific monoclonal antibody 3F8 in human melanoma cells. Oncoimmunology 4, e1023975, doi:10.1080/2162402X.2015.1023975 (2015).

36. Doronin, II et al. Ganglioside GD2 in reception and transduction of cell death signal in tumor cells. BMC Cancer 14, 295, doi:10.1186/1471-2407-14-295 (2014).

37. Gardai, S. J. et al. Cell-surface calreticulin initiates clearance of viable or apoptotic cells through trans-activation of LRP on the phagocyte. Cell 123, 321–334, doi:10.1016/j.cell.2005.08.032 (2005).

38. Lo, M. et al. Effector-attenuating Substitutions That Maintain Antibody Stability and Reduce Toxicity in Mice. J Biol Chem 292, 3900–3908, doi:10.1074/jbc.M116.767749 (2017).

39. Jaffe, N., Puri, A. & Gelderblom, H. Osteosarcoma: evolution of treatment paradigms. Sarcoma 2013, 203531, doi:10.1155/2013/203531 (2013).

40. Gibson, T. M. et al. Temporal patterns in the risk of chronic health conditions in survivors of childhood cancer diagnosed 1970-99: a report from the Childhood Cancer Survivor Study cohort. Lancet Oncol 19, 1590–1601, doi:10.1016/S1470-2045(18)30537-0 (2018).

41. Majzner, R. G. et al. Assessment of programmed death-ligand 1 expression and tumor-associated immune cells in pediatric cancer tissues. Cancer 123, 3807–3815, doi:10.1002/cncr.30724 (2017).

42. Aghighi, M. et al. Magnetic Resonance Imaging of Tumor-Associated Macrophages: Clinical Translation. Clin Cancer Res 24, 4110–4118, doi:10.1158/1078-0432.CCR-18-0673 (2018).

43. Ren, L. et al. Characterization of the metastatic phenotype of a panel of established osteosarcoma cells. Oncotarget 6, 29469–29481, doi:10.18632/oncotarget.5177 (2015).

44. Yoshida, S. et al. Ganglioside G(D2) in small cell lung cancer cell lines: enhancement of cell proliferation and mediation of apoptosis. Cancer Res 61, 4244–4252 (2001).

45. Iams, W. T., Porter, J. & Horn, L. Immunotherapeutic approaches for small-cell lung cancer. Nat Rev Clin Oncol 17, 300–312, doi:10.1038/s41571-019-0316-z (2020).

46. Rudin, C. M. et al. Pembrolizumab or Placebo Plus Etoposide and Platinum as First-Line Therapy for Extensive-Stage Small-Cell Lung Cancer: Randomized, Double-Blind, Phase III KEYNOTE-604 Study. J Clin Oncol 38, 2369–2379, doi:10.1200/JCO.20.00793 (2020).

47. Shan, D., Ledbetter, J. A. & Press, O. W. Signaling events involved in anti-CD20-induced apoptosis of malignant human B cells. Cancer Immunol Immunother 48, 673–683, doi:10.1007/s002620050016 (2000).

48. Chao, M. P. et al. Anti-CD47 antibody synergizes with rituximab to promote phagocytosis and eradicate non-Hodgkin lymphoma. Cell 142, 699–713, doi:10.1016/j.cell.2010.07.044 (2010).

49. Hudak, J. E., Canham, S. M. & Bertozzi, C. R. Glycocalyx engineering reveals a Siglec-based mechanism for NK cell immunoevasion. Nat Chem Biol 10, 69–75, doi:10.1038/nchembio.1388 (2014).

50. Wang, J. et al. Siglec-15 as an immune suppressor and potential target for normalization cancer immunotherapy. Nat Med 25, 656–666, doi:10.1038/s41591-019-0374-x (2019).

51. Siglec-15: An Attractive Immunotherapy Target. Cancer Discov 10, 7–8, doi:10.1158/2159-8290.CD-NB2019-136 (2020).

52. Majzner, R. G. et al. Tuning the Antigen Density Requirement for CAR T-cell Activity. Cancer Discov 10, 702–723, doi:10.1158/2159-8290.CD-19-0945 (2020).

53. Lynn, R. C. et al. c-Jun overexpression in CAR T cells induces exhaustion resistance. Nature 576, 293–300, doi:10.1038/s41586-019-1805-z (2019).

54. Weiskopf, K. et al. Engineered SIRPalpha variants as immunotherapeutic adjuvants to anticancer antibodies. Science 341, 88–91, doi:10.1126/science.1238856 (2013).

55. Gholamin, S. et al. Disrupting the CD47-SIRPalpha anti-phagocytic axis by a humanized anti-CD47 antibody is an efficacious treatment for malignant pediatric brain tumors. Sci Transl Med 9, doi:10.1126/scitranslmed.aaf2968 (2017).

56. Gray, M. A. et al. Targeted glycan degradation potentiates the anticancer immune response in vivo. Nat Chem Biol, doi:10.1038/s41589-020-0622-x (2020).

57. Wu, H. W. et al. Anti-CD105 Antibody Eliminates Tumor Microenvironment Cells and Enhances Anti-GD2 Antibody Immunotherapy of Neuroblastoma with Activated Natural Killer Cells. Clin Cancer Res 25, 4761–4774, doi:10.1158/1078-0432.CCR-18-3358 (2019).

